# Causal single-cell RNA-seq simulation, in silico perturbation, and GRN inference benchmarking using GRouNdGAN-Toolkit

**DOI:** 10.1101/2025.08.14.670294

**Authors:** Yazdan Zinati, Amin Emad

## Abstract

**Background:** Rapid advances in high-throughput single-cell sequencing technologies, coupled with the development of computational methods capable of leveraging large datasets, have led to the emergence of numerous approaches for deciphering regulatory interactions in the form of Gene Regulatory Networks (GRNs). However, in the absence of context-specific gold-standard ground truths, particularly those containing causal interactions, systematically benchmarking GRN inference methods remains a challenge. Thus, we previously developed GRouNdGAN, a causal implicit generative model capable of simulating realistic observational and interventional scRNA-seq data following a user-defined GRN on any biological system of interest. Importantly, we demonstrated that GRouNdGAN generates datasets that bridge the gap between experimentation and simulation for GRN inference benchmarking.

**Method:** Building upon the GRouNdGAN framework, we developed an extended toolkit that offers additional features, including interactive model visualization, training monitoring, more customizable GRN creation options, synthetic data similarity and GRN inference benchmarking metrics, and an intuitive TF knockout prediction module. Here, we provide a step-by-step procedure for implementing the protocol from start to finish and introduce alternative variations to adapt GRouNdGAN to studies with different experimental setups. GRouNdGAN-Toolkit is publicly available as a python code repository and containerized application and is accompanied by a tutorial website featuring a collection of simulated datasets. Model training largely depends on graphic hardware and the size and density of the input GRN, and typically takes around 75h to complete on a single GPU. Excluding model training, this protocol typically takes less than 25min to complete.

**Discussion:** GRouNdGAN-Toolkit is a versatile simulator with user friendly interface that does not assume advanced computational genomics expertise, enhancing its usability and accessibility across a wide range of users.

## Introduction and Background

Understanding how the gene expression is regulated within cells, including when, where, and to what extent a gene is transcribed and translated into a functional product like a protein, is fundamental to molecular biology[1, 2]. These regulatory processes, collectively known as gene regulation, are often conceptualized as transcriptional or gene regulatory networks (GRNs), which capture the dynamic interactions among genes and their regulators. To allow for computational analysis, GRNs are often modeled as directed acyclic bipartite graphs, linking transcription factors (TFs) to their target genes[3]. Inferring GRNs from bulk or single-cell RNA sequencing data[4–7] alone or in combination with other data modalities[8–10], has long been a topic of significant interest. This is due to the fact that GRNs play a critical role in our understanding of biological processes involved in cellular development, function, and disease, with applications in identifying therapeutic targets and predicting the outcomes of perturbation experiments[11]. However, systematically benchmarking GRN inference tools remains a challenge due to the absence of context-specific gold-standard ground truth networks and their associated datasets, particularly those that contain causal interactions[12]. While a ground truth GRN could be constructed from literature and curated biological databases of interactions, it may not accurately reflect the context- and cell type-specific regulatory programs of the biological system under study as they are aggregated across various datasets. Experimental validation through perturbation experiments could provide definitive evidence for the presence or absence of individual interactions, but such approaches are costly, time-consuming, and labor-intensive[11, 13, 14]. To address this, multiple GRN-guided simulators have been developed, offering an alternative means of generating benchmarking datasets[12, 15–17]. However, existing simulators often fail to impose regulatory relationships causally within their generated data and have key limitations (see our previous study for a detailed discussion[18]). Moreover, reported mismatches between benchmarking results obtained from experimental and simulated datasets raise concerns about their reliability[12].

### Development of the protocol

We developed GRouNdGAN[18], a generative deep learning model capable of simulating realistic single cell data, performing in-silico perturbation experiments, and systematically benchmarking GRN inference methods. GRouNdGAN combines and builds upon elements from previous studies on generative adversarial networks (GANs) namely scGAN[19] and causalGAN[20], to generate synthetic single cells mimicking those within a provided reference dataset while imposing causal relationships between TFs and genes as defined by a user-provided GRN.

Our approach follows a two-step training process. First, we train a Wasserstein GAN[21] with gradient penalty (WGAN-GP)[22] to generate realistic single cells without the regulatory constraints of the GRN. Once trained, we freeze the parameters of the generator to use as a “causal controller” in training a second WGAN-GP. In the second step, random Gaussian noise, along with TF expression values coming from the causal controller, are provided to a generator (called the “target generator”) which derives its neural network connections from the provided GRN. The masking of neural connections in the target generator ensures that target genes are causally expressed under the influence of their regulating TFs and some unobserved/unknown confounders (modelled by the inputted random noise unit). Both WGAN-GPs include a library size normalization layer as their last layer. Since only unnormalized TF expression values are needed from the causal controller, the LSN layer of the first WGAN-GP is discarded after it is trained in the first step. These TF expression values, along with the target gene expression values regenerated by the target generator, are then fed into the LSN layer of the target generator for final normalization.

Both WGAN-GPs are trained in an adversarial manner, with a “Critic” that evaluates generated samples by estimating the Wasserstein distance between the reference and simulated data distributions, and a generator that continuously refines its output to minimize this distance. Through this minimax game, both components co-evolve in a dynamic feedback loop, each improving in response to the other.

Unlike traditional GANs, GRouNdGAN employs two auxiliary networks, the labeler and anti-labeler, to ensure the target generator incorporates TF expression when generating target gene expression values, rather than relying solely on the input noise. These networks predict TF expression from target gene expression by minimizing the L2 loss between estimated and true TF values. The anti-labeler is trained exclusively on generated data, while the labeler is trained on both real and generated data to make further connection to the biology of the reference dataset. This approach reinforces causal TF-gene relationships within the GAN framework ensuring that regulatory consistency and realistic single-cell generation are optimized in tandem.

### Applications of the method

In our original study[18], we demonstrated that GRouNdGAN possesses essential properties for systematically benchmarking of GRN inference algorithms. This was achieved by generating benchmarking datasets based on six experimental datasets, and various causal GRN configurations. In summary, GRouNdGAN generates realistic scRNA-seq data while simultaneously imposing the causal relationships defined by the user (provided as an input GRN). It ensures that imposed edges are emphasized, making them recoverable by a high-performing GRN inference algorithm. Conversely, regulatory relationships that were intrinsic to the reference biological data, yet were not part of the input GRN are disrupted in the simulated dataset, limiting the potential of bias and ensuring a fair evaluation of GRN inference methods. We observed that benchmarking results of seven GRN inference methods using GRouNdGAN were consistent with insights previously found by BEELINE[12] from curated and experimental datasets, suggesting that GRouNdGAN effectively bridges the gap between experimental and simulated data benchmarks. Since then, GRouNdGAN has facilitated the development and evaluation of new GRN inference methods, such as LLM4GRN[3] and scRegulate[23], enabling researchers to carry out a systematic comparative evaluation of their method compared to the state-of-the-art. Figure 1 demonstrates an overview of the GRouNdGAN pipeline.

**Figure 1:**
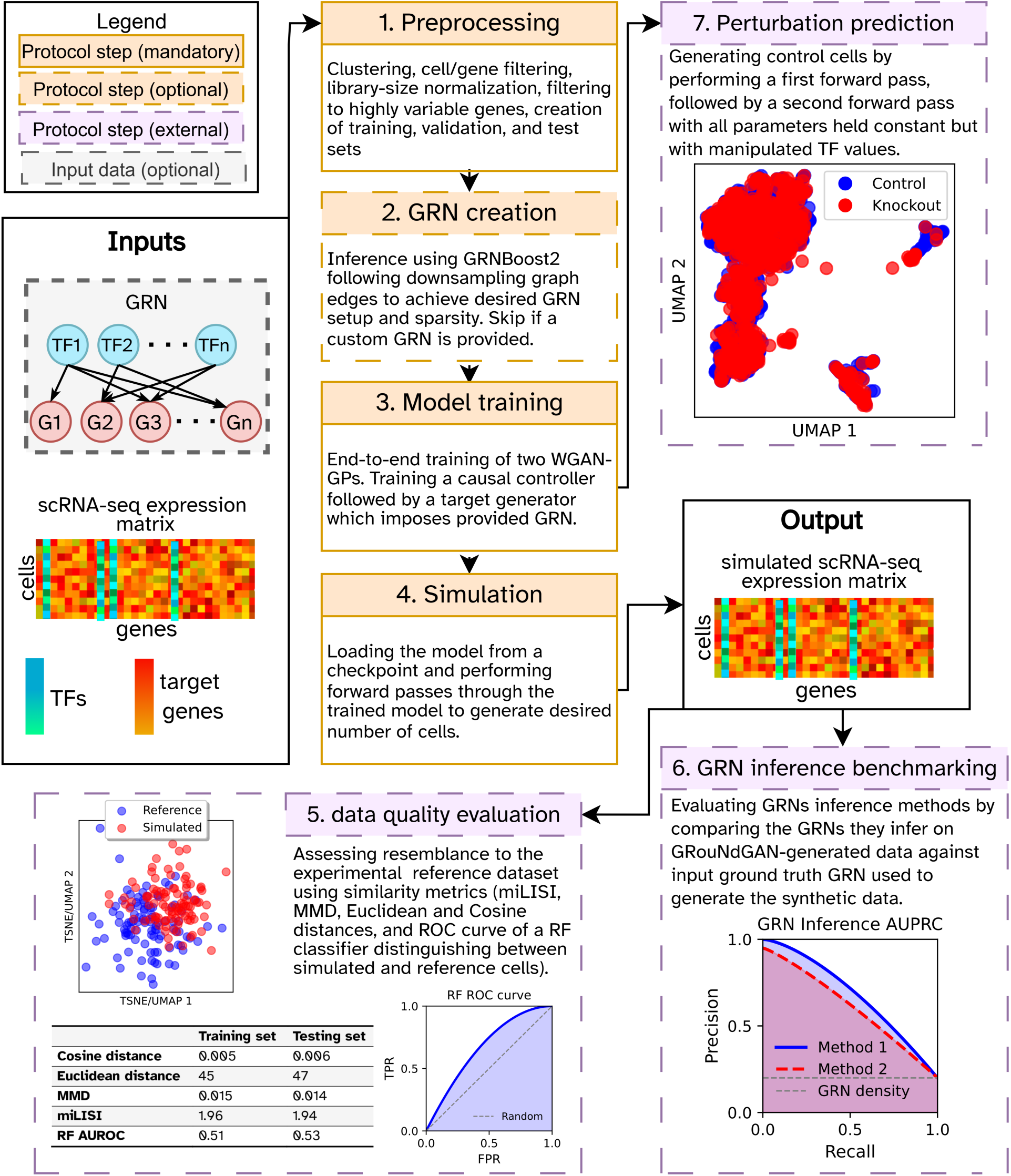
Schematic overview of the GRouNdGAN pipeline. Left: GRouNdGAN’s input data consists of a scRNA-seq dataset and an optional user-provided GRN. A list of TF names must be provided to differentiate TFs from target genes. Middle (light orange): Main steps in the GRouNdGAN protocol. The GRN creation step can be skipped only if a GRN is provided by the user. Right and bottom (in light purple): External protocol steps that involve using GRouNdGAN’s trained model or simulated data for secondary analysis and downstream tasks.

In addition to sampling from observational distributions to generate realistic benchmarking datasets, GRouNdGAN can also sample from interventional distributions and generate cells that do not naturally occur in the dataset, since it is a causal generative model. We previously showcased this application by performing in silico knock out experiments on top differentially expressed TFs of specific cell types, and observing their effect on the generated cells. The in silico perturbations can be performed during inference by manipulating the TF expressions generated by the causal controller, which are then fed to the target generator to produce the regulated target genes. In GRouNdGAN-Toolkit, we provide a user-friendly interface that allows users to generate cells under various perturbation conditions such as knockout. This application is not only suitable for predicting the outcome of specific perturbations but also for creating case/control experiments for studies that require such setups. Since our in silico perturbation is applied to the same batch of cells, keeping all generation parameters such as noise constant, it is deterministic, allowing to observe the effect of perturbations on the exact set of cells, providing matched case/control simulated datasets. This is not easily possible on experimental scRNA-seq data since most sequencing technologies are destructive, preventing the same cell from being sequenced twice (before and after perturbation).

As a specialized extension of general-purpose simulators, GRouNdGAN can fulfill all the same functions of non-causal simulators, when a GRN that accurately represents the reference dataset is imposed. Interestingly, we previously observed that general-purpose simulators that do not explicitly impose a causal graph, focus on generating realistic cells at the expense of compromising TF-gene relationships of the reference experimental dataset[18]. In this regard, GRouNdGAN may be a better choice even for tackling use cases typically handled by general-purpose simulators, as we have previously demonstrated that it outperforms state-of-the-art simulators in generating realistic data[18]. Here, we will briefly describe some use-cases of simulated data that GRouNdGAN can also support. Simulated data can be used for data augmentation to supplement limited sample availability caused by biological constraints, such as rare cell types, specific contexts, or populations. It can also help overcome technical limitations, like low-quality or low-yield sequencing data, and address ethical concerns related to human or animal experiments. The ability to generate synthetic data can significantly reduce the need for wet-lab experiments, which leads to considerable time and cost savings, particularly when testing biological hypotheses. Moreover, since GRouNdGAN can generate an unlimited number of cells without duplicates, it helps overcome challenges related to small sample sizes, which might not accurately reflect the broader population and can lead to poor robustness and reduced reproducibility of experimental results, though further validation is needed to confirm its effectiveness in this context.

In our original study, we showed that GRoundGAN-generated datasets effectively preserve key biological features such as cellular trajectories, pseudo-time ordering of cells, and marker gene expression. As a result, GRouNdGAN-generated data has the potential to improve downstream analyses, such as marker gene detection, classification of sparse cells, and trajectory inference, as previously demonstrated by scGAN[24] on their own data. Moreover, augmented data can enhance denoising and dropout imputation algorithms. Traditional imputation methods typically rely on borrowing information from similar cells within the dataset, but this approach can lead to oversmoothing, particularly for rarer cell populations, ultimately suppressing natural cell-to-cell variability. sciGAN[25] for instance, has demonstrated that synthetic cells can be used for information borrowing, rather than relying solely on observed cells, thus preserving the inherent stochasticity of the data. Finally, simulated data has the potential to be studied as means of sharing transcriptomic datasets at a reduced risk of exposing sensitive information, all while preserving the statistical and biological integrity of the original experimental reference data. However, demonstrating the effectiveness of this approach in addressing privacy concerns remains an ongoing challenge and is beyond the scope of our current work.

### Comparison with other methods

Several tools have been developed for simulation of scRNA-seq data over the years[24, 26–29]. However, most are general-purpose simulators that mainly focus on generating high-fidelity data that adheres to the statistical properties of experimental datasets. Using such simulators, one can augment data, improve downstream analysis, and benchmark some computational tools but cannot benchmark GRN inference algorithms. These simulators have shown to capture gene-gene correlations, but do not explicitly simulate data following a regulatory program that is either user-provided or extracted from the dataset, making them unsuitable for benchmarking GRN inference algorithms. Only a small number of simulators are explicitly guided by a user-defined GRN. One of the earliest models is GeneNetWeaver (GNW)[17] that was used to simulate bulk RNA-seq for benchmarking multiple GRN inference methods during Dialogue for Reverse Engineering Assessment and Methods (DREAM) challenges[17, 30, 31]. However, since it was originally designed for bulk RNA-seq data generation, its simulated datasets do not usually exhibit the statistical properties of single-cell data. Although there have been multiple attempts at adapting GNW to single-cell data, it falls short of modern simulators and cannot simulate data with multiple cell types, contexts, or trajectories[15].

SERGIO[15] and BoolODE[12] are single-cell GRN-guided simulators that use stochastic differential equations (SDEs) to model gene expression dynamics. BoolODE is a de novo simulator, meaning that it generates data entirely from scratch based solely on user-defined parameters and rules, without relying on a reference biological dataset. As a result, it cannot be used to benchmark GRN inference methods under specific contexts and conditions that are unique to an experimental dataset. Instead of a reference dataset, BoolODE relies on carefully selected user-defined parameters such as noise, dropout rate, and kinetic parameters to impose properties such as dropout, batch effect, variation across samples, population, and conditions. Although this gives the end-users greater control over the data generating process, it also places the responsibility of fully harnessing this flexibility on them, making it more challenging to generate new datasets. In other words, without extensive knowledge of single-cell transcriptomics and BoolODE’s inner workings, one may risk creating simulations that do not reflect experimental datasets. SERGIO[15] is a hybrid de novo/reference based simulator that first generates a “clean” dataset, strictly following a GRN without aligning to a reference dataset. The user then refines this simulated data by iteratively adjusting three parameters (controlling technical noise) to match a reference dataset, by assessing similarity based on five statistical measures: library sizes, mean gene expression, gene expression variance, and zero counts per gene and per cell. This approach introduces subjectivity that affects reproducibility of the results. In our experiments, even after the addition of technical noise, data quality remained subpar compared to other simulators. Furthermore, since technical noise is introduced after the causal relationships have already been encoded, it may unintentionally alter or disrupt interactions that should remain unchanged for a GRN benchmarking experiment. This can make relationships unidentifiable in the final dataset, potentially explaining the significant drop in the performance of all GRN inference methods tested by the authors in the original study[15].

On the other hand, GRouNdGAN optimizes its parameters to simultaneously impose causal TF-gene relationships while minimizing the distance between simulated and reference data distributions. This process is fully automated, requiring no user intervention or post-processing to preserve technical and biological noise, and cellular dynamics (such as pseudotime and trajectories). Unlike SERGIO and BoolODE, which generate pseudo-genes without clear mappings to the reference dataset, GRouNdGAN ensures a direct one-to-one correspondence between genes and TFs in both simulated and reference data. Each generated gene and TF mimic the expression patterns of their real counterparts. In contrast, preserving gene/TF identities in BoolODE and SERGIO requires manual fine-tuning of SDE parameters to align generated gene expressions with reference data, placing a significant burden on users. Finally, GRouNdGAN learns elaborate non-linear co-regulatory dynamics through implicitly parameterized functions implemented as neural networks from the data and without making simplifying assumptions. This is in contrast to the simplifying assumption of SERGIO that assumes the combinatorial effect of multiple regulators is the sum of their individual effects, or BoolODE that relies on a user-provided Boolean truth table for TF co-regulation rules.

### Experimental design

Besides configurations, GRouNdGAN requires only two inputs: (1) a scRNA-seq dataset (as the reference) and (2) a gene regulatory network (GRN). It produces a single output: a simulated scRNA-seq dataset that mimics the input reference dataset, while encoding the regulatory information from the input GRN within the simulated data. By preparing the inputs and using the output in a specific way (described in this manuscript), one can carry out different types of experiments, which we will explore in this section.

#### Input reference data and GRN considerations

GRouNdGAN’s architecture and optimization function ensures that data simulation is guided by a structural causal model defined by the input GRN. Unlike general-purpose simulators that prioritize generating realistic data often at the cost of losing underlying regulatory information, GRouNdGAN optimizes for both GRN imposition and the realism of the simulated data. One should however note that while GRouNdGAN can faithfully encode any GRN into the data it simulates, the resemblance of its output to the reference dataset deteriorates when the input GRN deviates considerably from the patterns present in the reference. In such cases, optimizing for both objectives (realistic data generation and GRN imposition) becomes contradictory rather than complementary. For this reason, we suggest users to impose GRNs that are generally in line with the reference dataset, though strict matching is not required for most applications. Having access to regulatory programs underlying the data is especially important for predicting the result of interventions. GRouNdGAN’s default approach involves building a scaffold GRN by inferring regulatory edges using GRNBoost2[5], then imposing a subset by retaining the top k interactions for each gene. The value of k can be adjusted to control the sparsity of the graph, with a denser GRN typically providing more information and leading to higher-quality simulations.

Instead of the default approach, one can use a custom GRN. Custom GRNs can be constructed by selecting a subset of edges other than top k (for example top 1^st^, 3^rd^, 5^th^, …) or those inferred by an algorithm other than the default option. Additionally, they can be constructed using data sourced from literature, structured curated biological databases (e.g., KEEG[32], I2D[33], IID[34], ESCAPE[35], and GTRD[36]) or other modalities(e.g., TF ChIP-seq or ATAC-seq). These sources can also be combined to create a more complete GRN. The edges in the input GRN do not have to reflect the true causal relationships in the reference dataset. They could arise from spurious correlations or other observed associations. However, once imposed by GRouNdGAN, these edges become the underlying causal edges of the simulated dataset.

In our original study[18], we imposed a single shared GRN on a highly heterogeneous dataset of human peripheral blood mononuclear cells (PBMC) and obtained state-of-the-art performance. However, since regulatory programs are cell type- and context-specific, it may be beneficial to impose different GRNs on subpopulations of the dataset, rather than using a single shared GRN, depending on the application. This can be easily achieved by training multiple population-specific GRouNdGAN models, each using a subpopulation of the experimental dataset along with a GRN specific to that subpopulation. To demonstrate this experimental setup, here we trained five GRouNdGAN models using cell-type-specific GRNs for the five most populated cell types in the PBMC dataset[37]. We followed the procedure outlined in Figure 2-a where preprocessing is done on the entire dataset after which it is separated based on cell type and finally used to construct cell type-specific GRNs. These cell type-specific GRNs depict distinct regulatory programs as shown by Figure 2-b, while still sharing some regulatory edges. Table 1 highlights strong performance of cell type-specific models across five metrics in generating realistic data. While this approach offers some advantages, it cannot leverage the potential benefits of information sharing, nor the performance improvements that come from using a larger training sample. Nonetheless, the cell type-specific models still achieved performance comparable to that of the non-cell-type specific model (PBMC-ALL in Table 1) despite being trained on fewer samples (See Table S1). By aggregating the outputs from cell type-specific models in proportion to the relative abundance of each cell type, one can effectively reconstruct the overall dataset simulation, similar to a non-cell-type specific model (see Figure 2-c).

**Figure 2:**
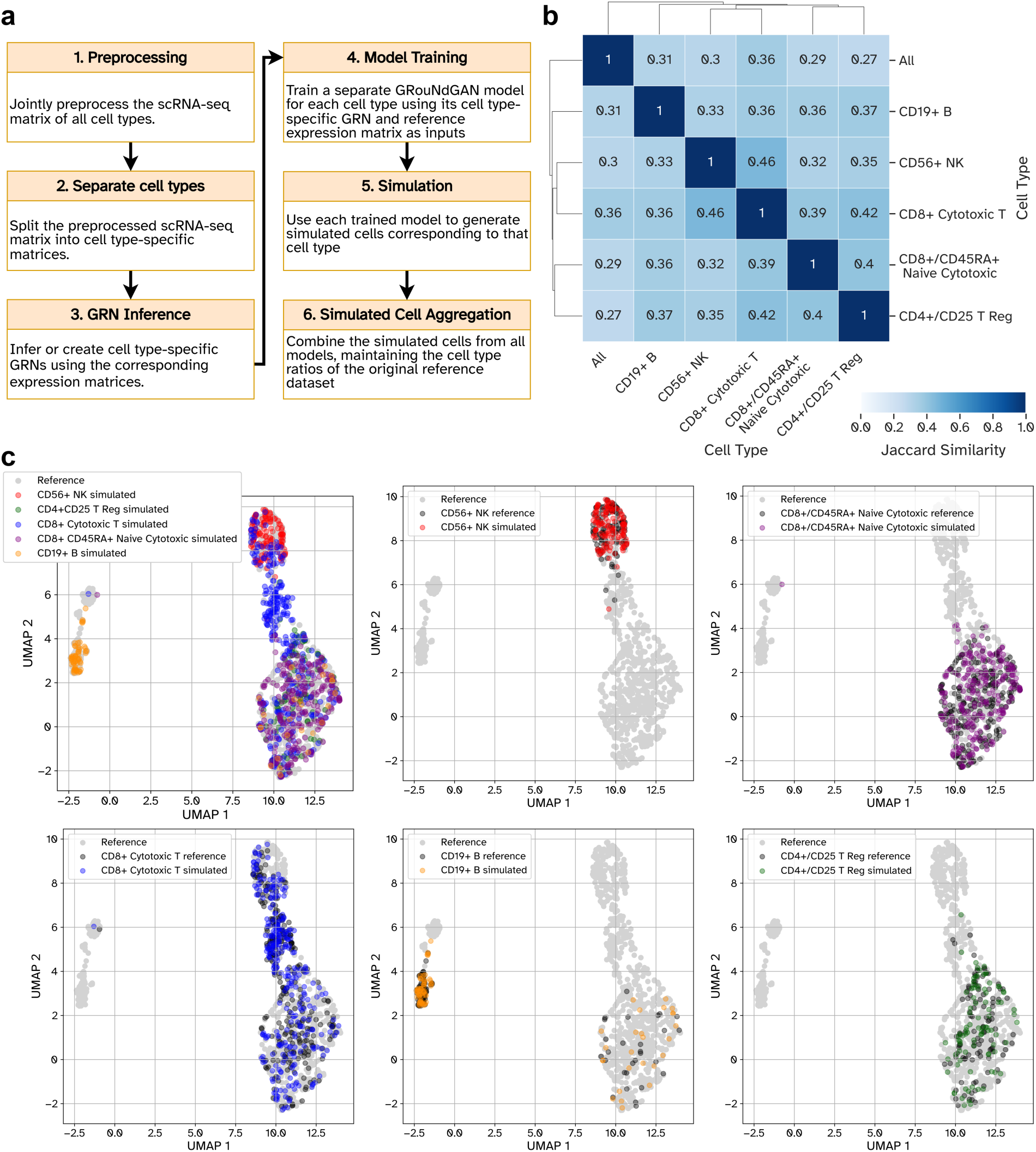
Cell type-specific simulation of cells using GRouNdGAN. Panel **a** illustrates an experimental setup designed for running simulations with cell type-specific GRNs. **b** Clustered heatmap showing pairwise Jaccard similarity between GRNs constructed for each cell type, based on shared TF-gene regulatory edges. Hierarchical clustering was done using average linkage and Euclidean distance. **c** Subplots show the UMAP embedding of cell type-specific simulations overlaid on reference (real) cells. The top left subfigure displays an aggregated UMAP where simulated cells from all cell type-specific models are combined in proportions matching the reference dataset (see Table S1 for relative abundance of cell types). The overlaid reference cells serve as a visual benchmark for simulation data quality.

**Table 1:**
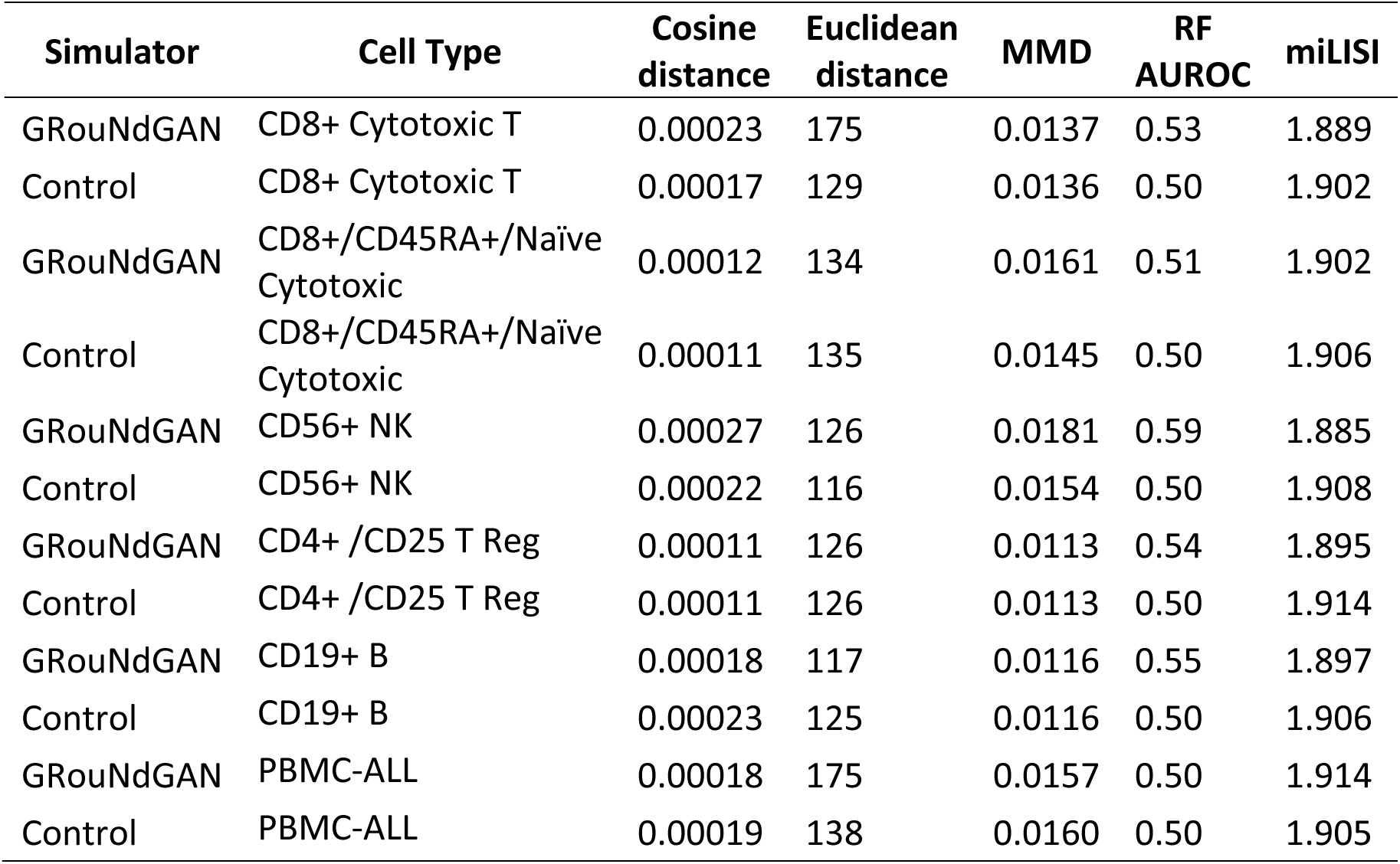
Performance of GRouNdGAN in generating realistic scRNA-seq data using cell-type-specific GRNs using the PBMC dataset. The metrics are calculated by comparing a held-out test set of 1000 real cells and a simulated dataset of the same size. In the imposed GRN each gene is regulated by a maximum of 15 TFs. The default algorithm GRNBoost2 was used with the training experimental data to form the imposed GRN. For the cosine distance, Euclidean distance, and MMD, values closer to zero are desired. For RF AUROC a value closer to 0.5 and for miLISI a value clo<u>ser to 2 reflect a more realistic simulation.</u>

When there is no need to integrate the simulation across cell types, the dataset may first be partitioned by cell type, followed by independent preprocessing and GRN construction for each subset. This approach may offer some advantages: for example, one can separately preprocess and identify highly variable genes specific to each cell type. As a result, selected genes and TFs are more biologically relevant to specific cell types, leading to the inference and construction of more representative GRNs. Table S2 summarizes the performance of cell type-specific models under this alternative experimental setup.

For comprehensive benchmarking studies, we recommend that users generate multiple simulated datasets with varying characteristics such as different GRN densities, different topologies, different number of TFs, and different number of target genes. Another approach that can enhance the rigor of a benchmarking study (previously demonstrated in our original study) involves creating two distinct GRNs. Each GRN would consist of half of the top k regulating TFs for each gene, with one serving as an unimposed control and the other used for simulation. The unimposed control GRN can serve as baseline for computing performance metrics.

#### Output data considerations

GRouNdGAN includes built-in functionalities to assess both data quality and GRN inference performance. Data quality is evaluated using the Area Under the Receiver Operating Characteristic Curve (AUROC) of a Random Forest (RF) distinguishing experimental from simulated cells, mean integration local inverse Simpson’s index (miLISI)[38], Maximum Mean Discrepancy (MMD), and the Euclidean and Cosine distances between the centroids of experimental and simulated cells. GRN inference performance is measured with the Area Under the Precision-Recall Curve (AUPRC) and precision. For detailed explanations of each metric, please refer to our original publication.

A key advantage of using simulation for benchmarking GRN inference methods is the ability to generate an unlimited number of samples, which offers significant benefits over subsampling a finite experimental dataset. This enables comprehensive benchmarking by assessing GRN inference algorithms in terms of computational resource usage (e.g., time and memory efficiency on large datasets), and enables quantifying their stability across multiple runs, evaluating their robustness to noise, and optimizing their performance for specific applications. To demonstrate this, we assessed the performance of PIDC[6] and PPCOR[7] on simulated datasets ranging from 100 to 100,000 cells, with the results presented in Figure 3. The reference dataset (PBMC-CTL) for this consists of the most abundant cell type of the PBMC dataset[37] (i.e., CD8+ Cytotoxic T cells). Figure 3-a, b, c show that GRouNdGAN-generated cells closely resemble real cells across multiple quality metrics and in low-dimensional t-SNE space. These results are based on 100 replicate comparisons, comparing a held-out test of 1000 real cells and 100 sets of simulated cells (each containing 1000 cells for a total of 100,000), demonstrating consistent and high data quality of GRouNdGAN simulations.

**Figure 3:**
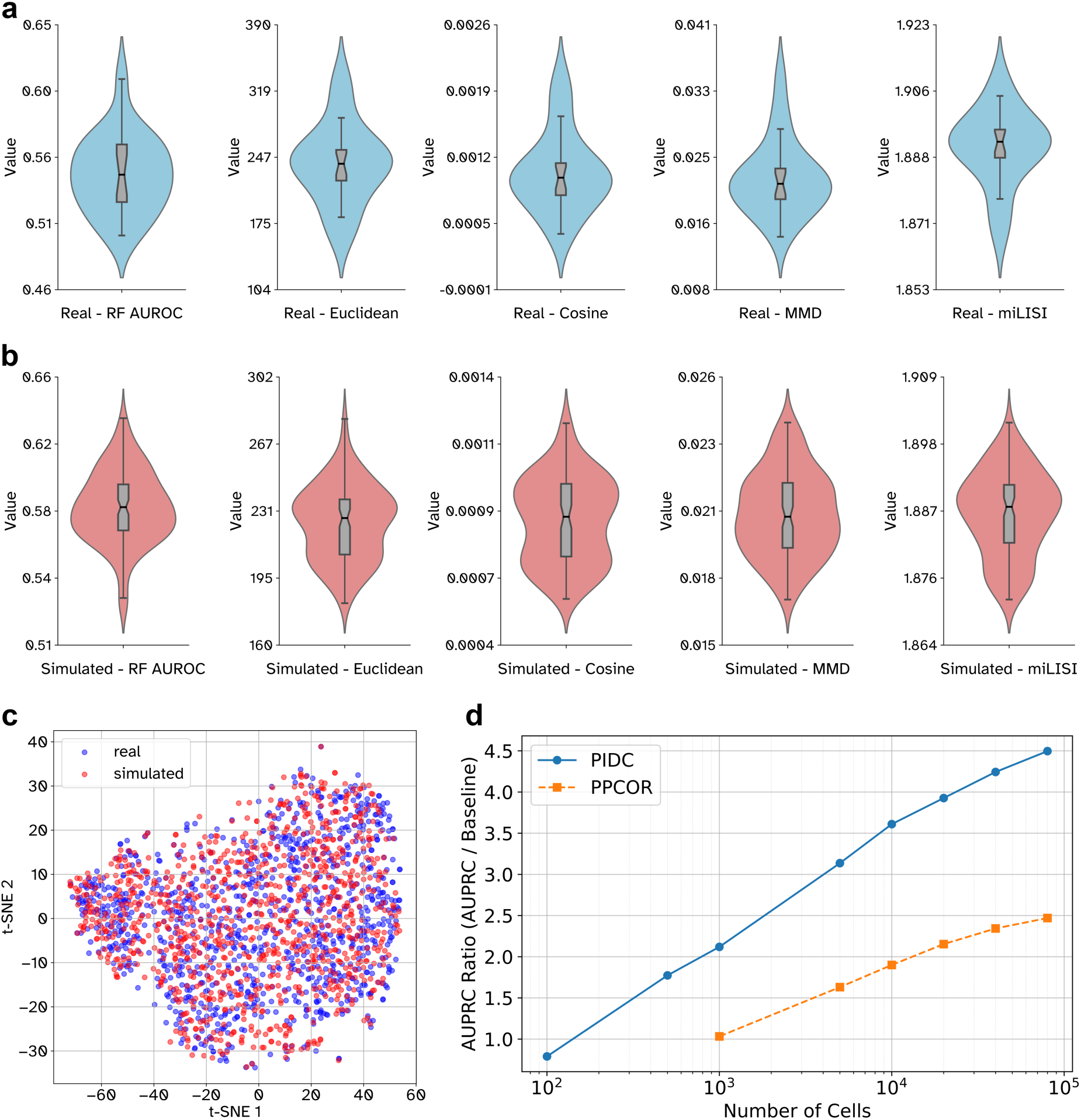
Robustness analysis of simulated data. **a** Violin plots showing five data quality metrics computed between partitions of 19000 reference training experimental cells (1000 cells each) and a held-out test set of 1000 experimental PBMC-CTL cells, serving as real vs real control benchmark. **b** Similar to **a** but computed between 100 distinct simulated datasets (each containing 1000 cells) and the same experimental test set. **c** t-SNE plot of 1000 simulated cells and the same 1000-cell test showing a visual assessment of the distributional similarity between generated and experimental cells. **d** AUPRC ratio (compared against the GRN density) for PIDC and PPCOR in inferring GRNs from simulated GRouNdGAN scRNA-seq data. The x-axis (log scale) represents the number of cells provided. The imposed GRN contains the top 15 regulating TFs for each target gene. Performance improves when increasing the number of provided cells, with PIDC outperforming PPCOR. PIDC takes over 12 hours to run on more than 80000 cells and PPCOR fails to output a GRN when provided with 100 and 500 cells.

As expected, we observed an improvement in AUPRC for GRN recovery as the number of cells increased (Figure 3-d) with PIDC outperforming PPCOR. PPCOR fails on very small datasets (100– 500 cells) and PIDC becomes computationally prohibitive (taking longer than 12 hours to run) on large datasets (>80,000 cells). These results underscore that GRouNdGAN can not only be used for data augmentation to increase the stability of downstream computational tools, but also to systematically evaluate their robustness and scalability.

### Expertise needed to implement the protocol

Some familiarity with the command line and basic understanding of the Python programming language are required to execute this protocol. Our protocol assumes no advanced bioinformatics expertise; though, such expertise may be necessary for downstream tasks and analysis involving the output of this protocol.

### Discussion and Limitations

While GRouNdGAN is a powerful generative model, it has inherent limitations that need to be considered. Through adversarial training, GRouNdGAN learns to approximate the distribution of training samples. Once trained, it samples simulated cells from the learned probability distribution. As such, like most reference-based simulators, its ability to generalize and produce realistic outputs depends not only on the number but also the quality and diversity of training examples. Despite that, our previous study revealed that GRouNdGAN can introduce causality and capture properties such as pseudo-time ordering and cellular trajectories on relatively small datasets and excessively large datasets do not substantially improve performance beyond a certain point.

Throughout all our analyses, we consistently used the same model architecture and hyperparameter settings, achieving state-of-the-art performance with different choices of GRN across diverse datasets spanning various cell-types, conditions, species, and sequencing technologies. However, when training GRouNdGAN to a completely new dataset, especially one with a significantly different number of genes or transcription factors, we recommend fine-tuning hyperparameters using a validation set. GRouNdGAN’s current architecture incorporates a library-size normalization (LSN) function as the last layer of its causal generator. The LSN layer ensures that all generated cells maintain a consistent fixed library size across selected genes and TFs. Our ablation study[18] confirmed that this layer strengthens GRN imposition and enhances the quality of generated cells. Additionally, our observations indicate that it accelerates model convergence and increases training stability, findings that align with previous GAN-based research in scRNA-seq[24]. While we offer the option to use GRouNdGAN without the LSN layer for flexibility in specific biological analyses requiring unnormalized count data, we advise against this unless necessary as it reduces the quality of the simulated data.

GRouNdGAN’s main objective is two-fold: to generate realistic scRNA-seq data that aligns with the reference dataset and to impose causal relationships of a given GRN. If the TF-gene regulatory relationships described by the GRN are overly inconsistent with the patterns in the reference data, the two objectives become contradictory and impossible to optimize simultaneously. Conversely, providing a GRN consistent with the reference dataset can serve as additional knowledge, improving the quality of simulated cells compared to simulators that do not incorporate regulatory information. Hence, we suggest creating the input GRN from the dataset itself or using other modalities pertaining to the same biological system under study.

The model is currently confined to capturing gene regulation through a bipartite GRN (TF to gene). Gene regulation is inherently more complex, involving gene-gene interactions, multi-level regulation, feedback loops, and self-regulation—elements this model doesn’t fully capture.

However, we should mention that this simplified GRN model is widely used by GRN inference algorithms alike and avoids overcomplicating computations. We plan to relax this limitation as much as possible in future versions of GRouNdGAN.

## Materials and Methods

### Equipment

#### Hardware

This protocol was run on a high-performance compute cluster with 2 cores of an AMD EPYC 7413 (@ 2.65 GHz, 128M cache L3) processor, 16GB of RAM, and a single NVidia A100SXM4 (40 GB memory) GPU. Additionally, we conducted tests of the protocol on a desktop workstation equipped with an 11th Gen Intel Core i7-11700 processor (@ 2.50 GHz, 8 cores), 32GB of RAM, and a GeForce RTX 3070 GPU (24GB memory). At minimum, we recommend a dual-core computer with 16GB of RAM, and a GPU with 24GB of video memory. However, it is still possible to run GRouNdGAN on devices with smaller VRAM than 24GB by making the input GRN sparser or reducing the model size through hyperparameters. An internet connection is required for setting up the compute environment and downloading GRouNdGAN and input data files.

#### Software

Operating system: Linux, Windows, or MacOS

- Python version 3.9.6
- Docker or Singularity if you chose to run GRouNdGAN via containerized environments.
- GRouNdGAN: A free and open-source implementation is maintained on GitHub at https://github.com/Emad-COMBINE-lab/GRouNdGAN.
- basic commands such as curl, tar, git, etc.
- (Optional) Jupyter’s interactive computing environment for secondary analysis
- (Optional) pyenv to install and switch between GRouNdGAN’s required Python version and other installed versions on the system and its pyenv-virtualenv plugin to manage python virtual environments.
- (Optional) SCANPY[39], Seurat[40], or other single-cell processing pipelines for converting raw single-cell expression data into appropriate input formats.

#### Configuration file

Each GRouNdGAN command requires providing the path to a configuration file which specifies the protocol’s hyperparameters and inputs. This configuration file (with a `.cfg` extension) employs a structure closely similar to that of Microsoft Windows INI files, which can be easily customized by the users. Aside from configurations regarding input and output parameters (discussed in each procedure step), the content of the base configuration file can remain unchanged. GRouNdGAN has consistently well-performed across datasets with the same set of hyperparameters specified in the base configuration file. Although GRouNdGAN is not overly sensitive to the choice of hyperparameters, we still advise to test different hyperparameters on a validation set when training on a new dataset with significantly different dimensions in terms of number of TFs or target genes. A default base configuration file is available in Supplementary Notes and our code repository (path: `configs/causal_gan.cfg`) but can also be directly downloaded from the command line by running the following command:

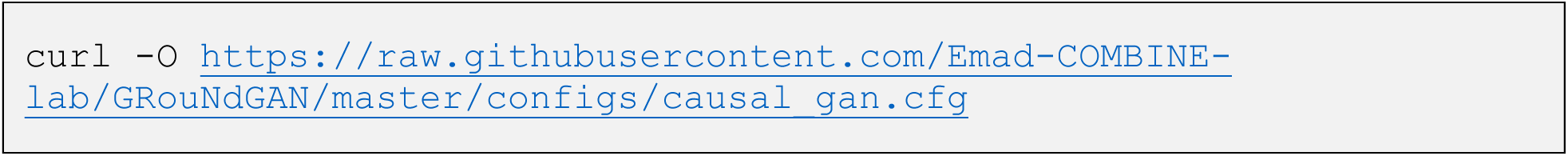

#### Input data files

The protocol requires two user data inputs: raw single-cell RNA-seq count data and a list of known TFs belonging to the species being studied. GRouNdGAN accepts the scRNA-seq matrix as an AnnData object stored in the standard hierarchical `.h5ad` format. The AnnData object, at a minimum, requires the `X` and `var` fields to contain the expression data matrix and gene identifiers (e.g., Ensembl Gene ID, HGNC (HUGO Gene Nomenclature Committee) Symbol, or NCBI Entrez Gene ID), respectively. Note that the expression matrix needs to be stored as a dense (and not sparse) matrix. Alternatively, GRouNdGAN can accept 10x Genomics-formatted directories that include `matrix.mtx` (gene expression matrix in Matrix Market Exchange format), `genes.tsv` (list of features to be used as gene identifiers as discussed), and `barcodes.tsv` (list of cell barcodes representing unique cells) files. Users can optionally supplement that input with a `.tsv` file for additional cellular annotations and metadata such as cell type and cluster. To match TFs to their expression data, the list of known TFs must use the same identifier type that were chosen to uniquely name genes in the count matrix. Users can decide to omit GRouNdGAN’s default causal GRN preparation procedure and instead define a custom causal graph. In that case, a causal graph should be supplied in lieu of the TF list. See Procedure – Step 2 for the causal graph’s format.

### Equipment setup

#### Software Installation

Depending on user preferences and requirements, GRouNdGAN can either be locally installed or run as a containerized application. Local installation is recommended for users that need greater control over GRouNdGAN’s environment or decide to build new projects upon its framework as a foundation for new bioinformatics work. To reproduce our results or run the protocol using a pre-configured environment where installation is not needed, we suggest using our Docker or Singularity (where Docker is not allowed; e.g, HPC clusters) images, both of which are configured with CUDA support for GPU acceleration. GRouNdGAN’s latest version can be downloaded from our GitHub repository’s release page: https://github.com/Emad-COMBINE-lab/GRouNdGAN/releases/.

##### Local Installation

###### TIMING ∼12 min

For local installation, python 3.9.6 is required. We recommend isolating GRouNdGAN’s environment from the base environment to avoid breaking package dependencies either using python’s virtualenv module or pyenv coupled with pyenv-virtualenv to manage multiple python versions. Here, we show how to set up GRouNdGAN with Python’s own virtual environment module (venv) and package management system (pip).

1. Start with cloning GRouNdGAN’s GitHub repository to a directory of your choosing.

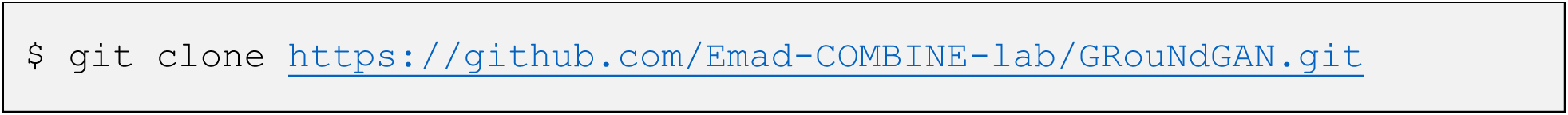

To fully reproduce our primary study[41], one can additionally include specific versions of scGAN, BEELINE[12], scDESIGN2[42], and SPARSIM[26] projects as submodules when cloning the repository.

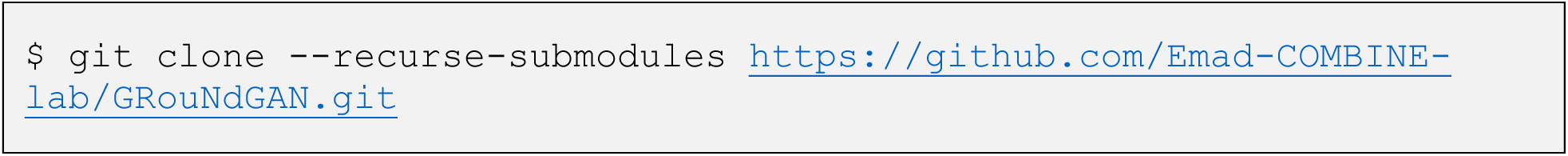

2. Navigate to the cloned directory.

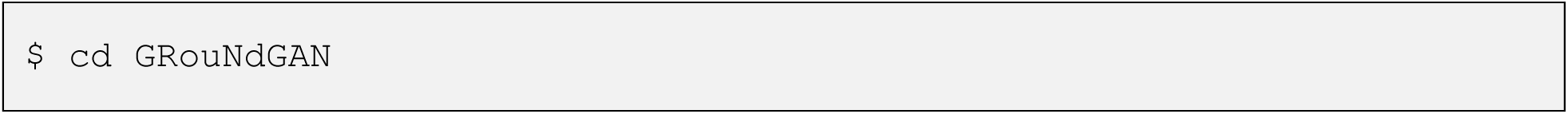

3. Initialize a new virtual environment for the project (here named `groundgan_env`).

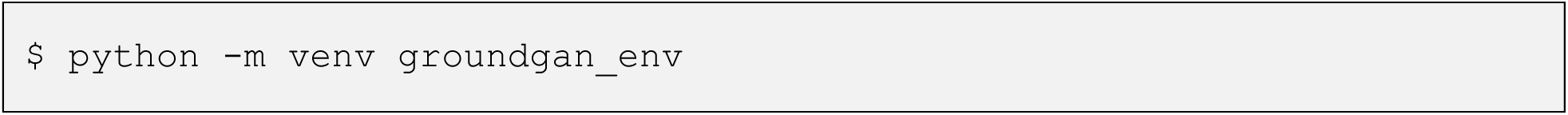

If you chose to use pyenv and pyenv-virtualenv, visit their tutorial and installation guides for your operating system at https://github.com/pyenv/pyenv and https://github.com/pyenv/pyenv-virtualenv.

4. Activate the created virtual environment.

Linus or MacOS:

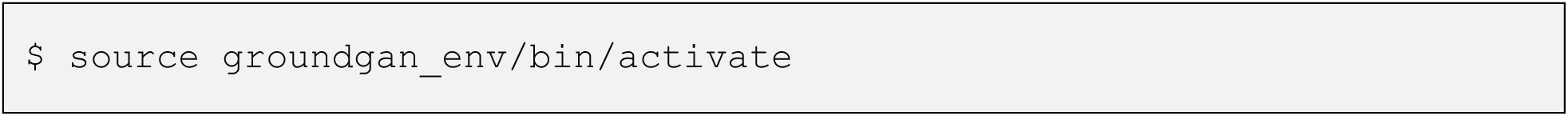

Windows:

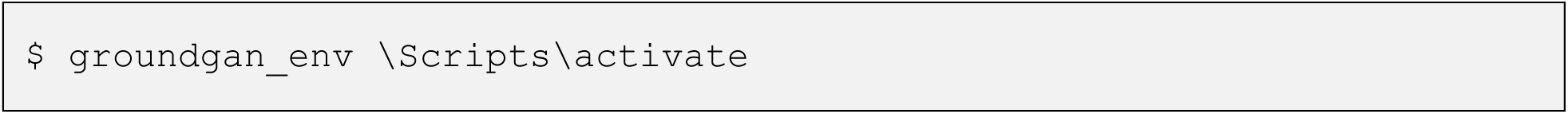

5. Project dependencies can now be installed in the created virtual environment through pip from the Python package index (PyPi) using the `requirements.txt` file available at the root of the cloned directory.

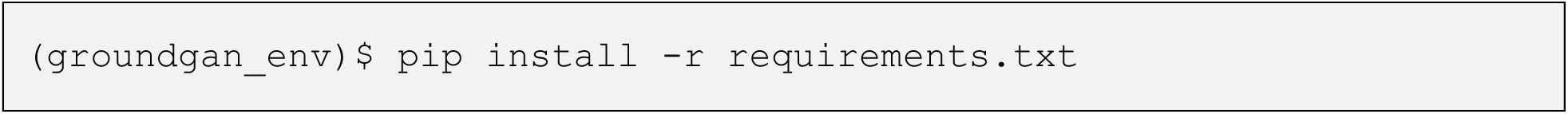

##### Using the Docker Image

###### TIMING ∼6 min

GRouNdGAN’s docker image is available on the DockerHub at https://hub.docker.com/r/yazdanz/groundgan.

1. Run the following command to pull the latest image from DockerHub:

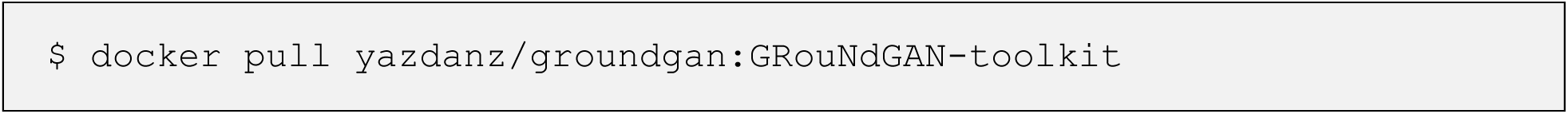

2. To run the Docker container, execute the following:

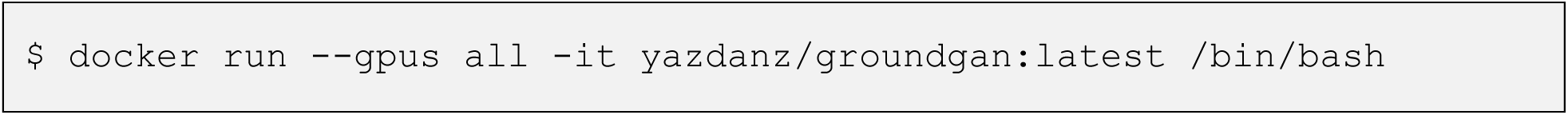

The –-gpus all and –it flags enable GPU support within the container and start an interactive terminal session respectively.

##### Building Docker Image from Dockerfile

###### TIMING ∼13 min

We only recommend this option if you wish to further finetune the provided containerized environment to your specific requirements and maintain reproducibility.

1. Clone GRouNdGAN’s GitHub repository to a directory of your choosing.

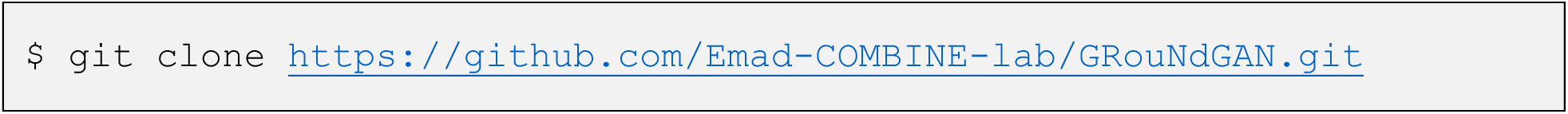

2. Navigate to the project directory.

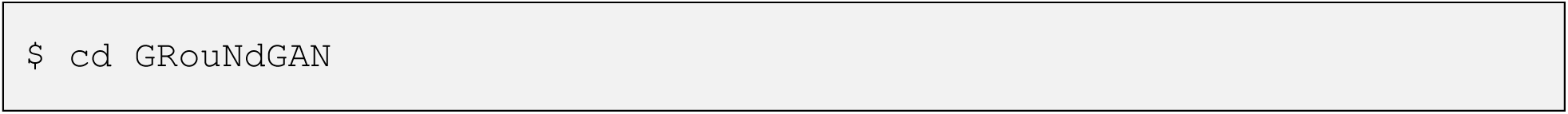

3. Make intended updates and modifications to the Dockerfile (path: `docker/Dockerfile`)

4. Build the Docker image using the provided Dockerfile. The command below will build a docker image with the tag `yourusername/groundgan:custom` which you can change.

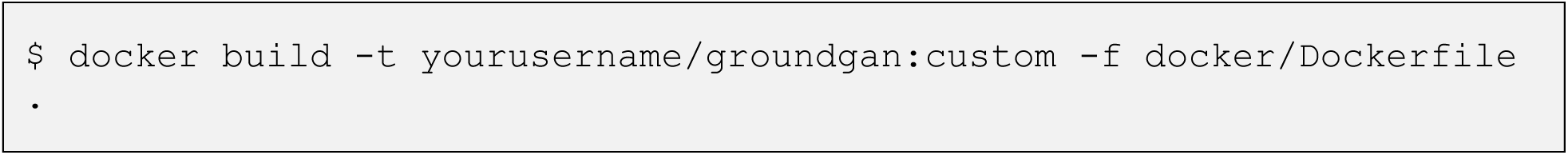

5. Once the image is built, start the docker container interactively and pass GPU devices with the following command. You can also decide to publish it to DockerHub.

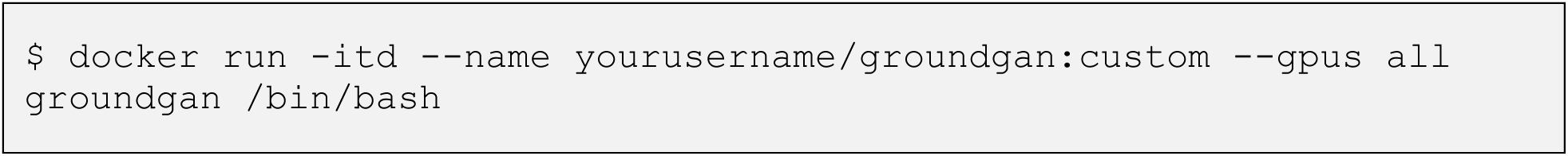

##### Using Singularity

###### TIMING ∼7 min

1. Pull and convert the container image from Dockerhub into the Singularity Image Format (SIF).

The following command creates a singularity image with the groundgan.sif filename.

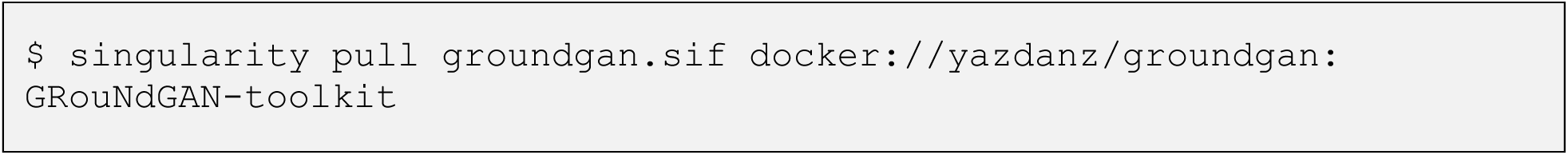

2. Start an interactive shell session within the `groundgan.sif` container. The `--nv` flag enables NVIDIA GPU hardware support and exposes the host system’s drivers and libraries to the container.

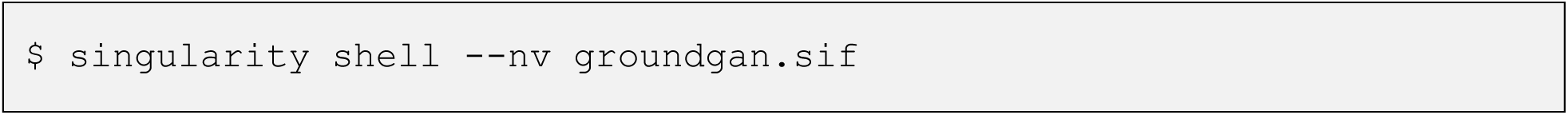

You can now execute shell commands inside the running container.

##### Verifying GPU Acceleration

###### TIMING <1 s

You can ensure that your device’s GPUs are recognized inside Docker and Singularity containers by running the `nvidia-smi` command. You should see detailed information regarding your GPU and installed drivers. See an example of the expected output below.

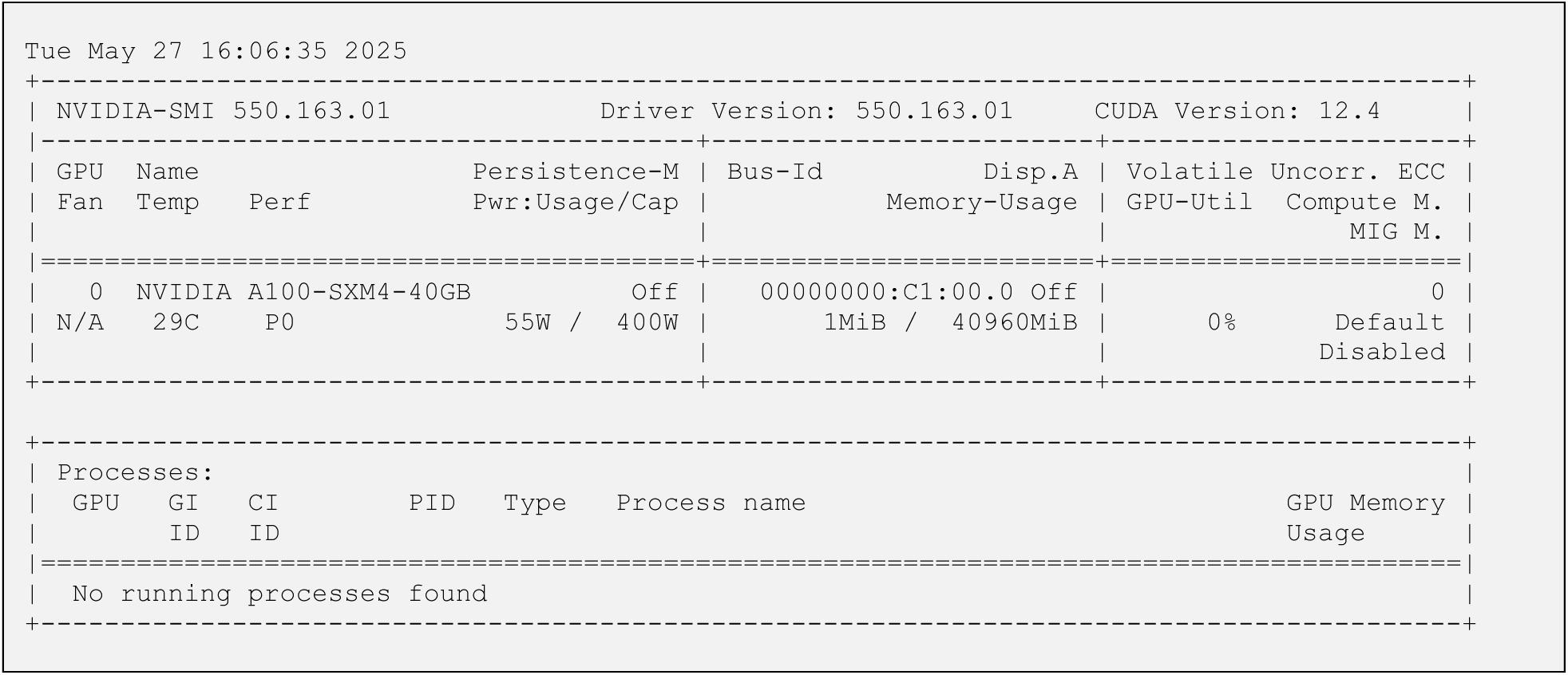

#### Example input data

Example input data comprising of the PBMC[37] and BoneMarrow[43] expression data and list of Homo sapiens and Mus musculus TFs (from AnimalTFDB3.0[44]) can downloaded from our webserver at https://nextcloud.computecanada.ca/index.php/s/pXKQ2isr47AwKEX. Alternatively, one can run the below command in the terminal to download, extract, and copy the raw input data files in the `data/raw` project directory. Make sure to run this command in the GRouNdGAN project directory after having cloned the repository.

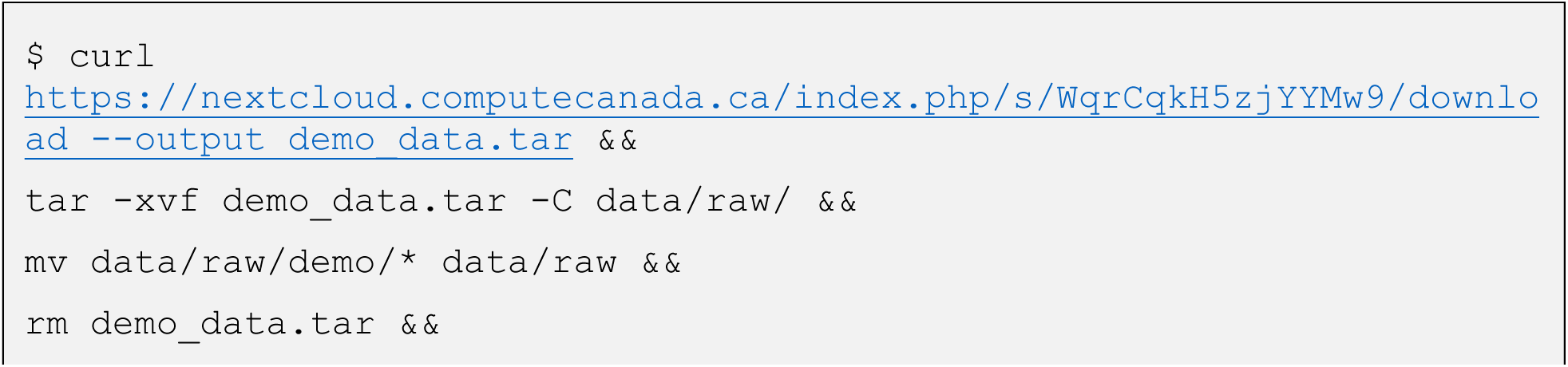
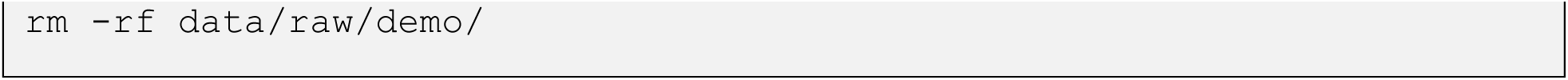

▴ **CRITICAL** Docker and Singularity images come prepackaged with the said example input data files and this step should be skipped.

## Procedure

GRouNdGAN can be executed through a command-line interface (CLI) by running the `main.py` file in the `src/` directory via the Python interpreter. Running `main.py` with the `--help` (or the abbreviated `-h`) flag outputs a summary of the tool and a list of required and optional arguments. Commands are listed as optional arguments and the path to the configuration file, which needs to be inputted alongside each command, is a required argument. Each of the seven available commands is a step in the procedure: `—preprocess`, `--create_grn`, `--train`, `--generate`,

`--evaluate`, `--benchmark_grn`, and `–perturb`. Below is the output of the `–-help` command.

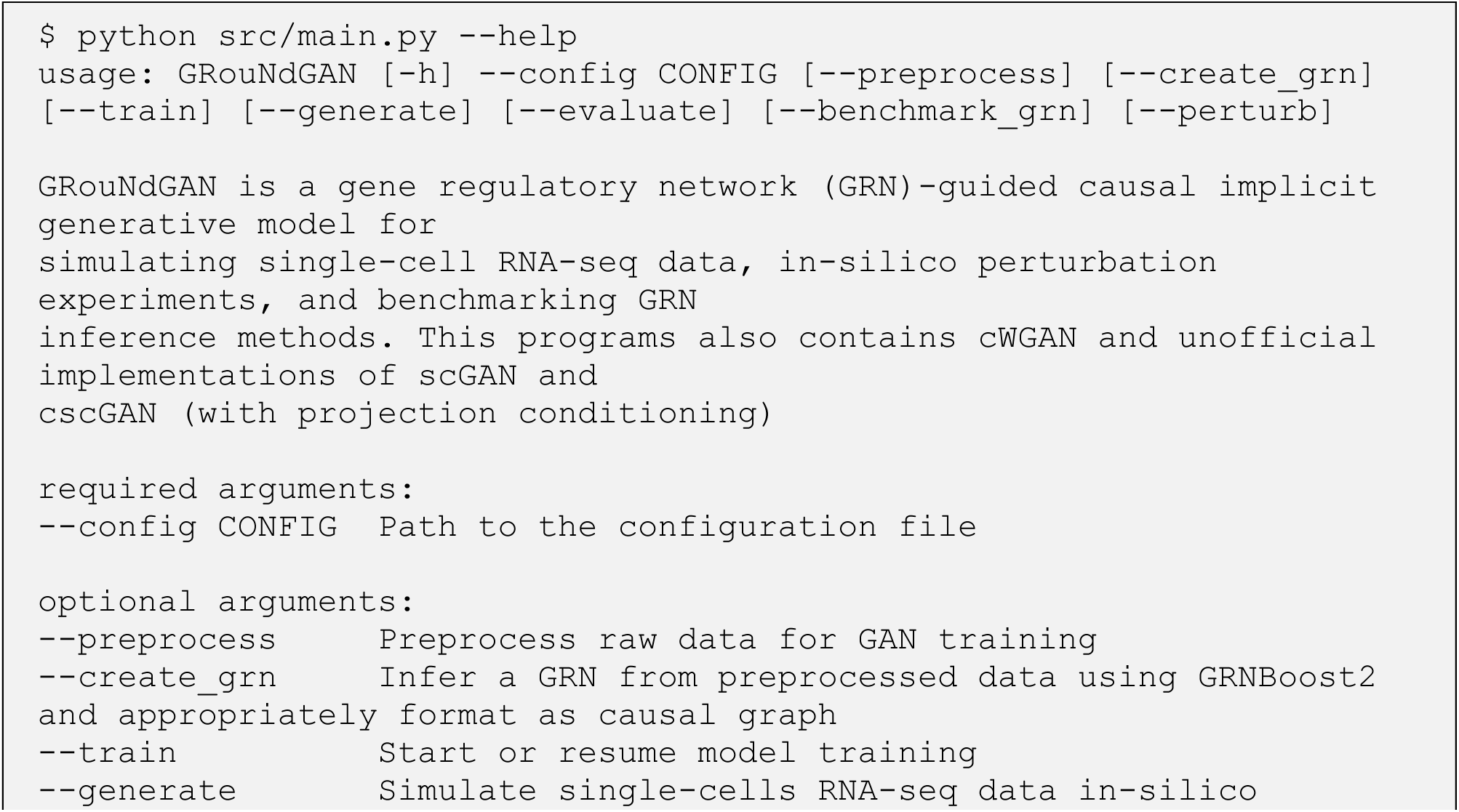
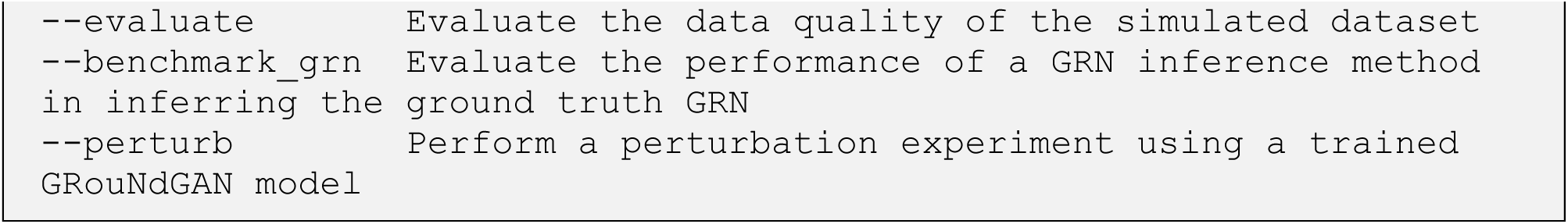

It is possible to execute the whole pipeline in one go through a single command by chaining multiple steps of the pipeline, as shown below.

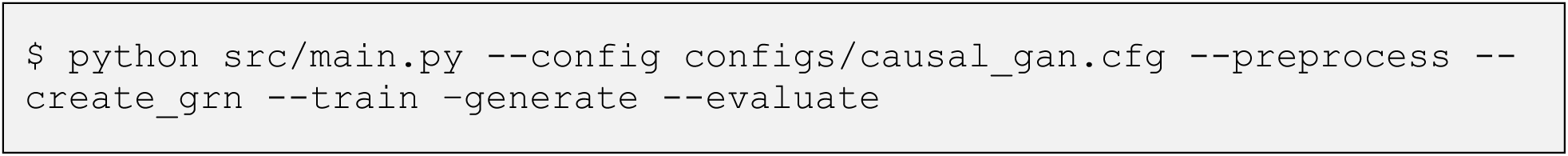

Alternatively, one can run each step separately. For example, the below command only runs the preprocessing step.

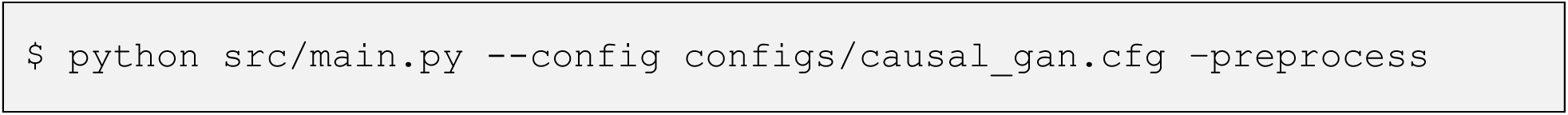

▴ **CRITICAL** If you decide to chain two or more commands, the supplied configuration file must include required arguments of all invoked commands. We discuss each step of the pipeline and relevant configuration file arguments in the following sections.

▲ **CRITICAL** Avoid chaining commands that are not meant to be run consecutively (e.g., running

--preprocess with --generate since this jumps over the --create_GRN and –train commands).

▴ **CRITICAL** Substitute python with python3.9 if you are running GRouNdGAN in a Docker or Singularity container.

### Step 1. Preprocessing

#### TIMING <1 min

Before running the preprocessing command, ensure that the configuration file contains the required arguments to the Preprocessing and Data sections of the configuration file, as outlined in Table 2.

**Table 2:** Arguments relevant to the preprocessing command in the configuration file.

| Configuration File Section | Argument | Explanation |
| --- | --- | --- |
| [EXPERIMENT]/[Preprocessing] | 10x<br>[type: boolean] | “True” if you are supplying 10x-formatted data files (`matrix.mtx`, `genes.tsv`, and `barcodes.tsv`).<br>“False” if you are providing annotated data (`anndata`) objects in the `.h5ad` format. |
| [EXPERIMENT]/[Preprocessing] | Raw<br>[type: file path/directory] | If 10x is set to “True”, path to the directory containing 10x-formatted data files.<br>Otherwise, path to the raw (unprocessed) `.h5ad` file. |
| [EXPERIMENT]/[Preprocessing] | validation set size<br>[type: integer] | Number of cells to include in the validation set. |
| [EXPERIMENT]/[Preprocessing] | test set size<br>[type: integer] | Number of cells to include in the test set. |
| [EXPERIMENT]/[Preprocessing] | Annotations<br>[type: directory, Optional] | Path to additional annotations (such as cell-type, clusters, etc. for cells/samples) |
| [EXPERIMENT]/[Preprocessing] | min cells<br>[type: integer] | Filter genes that are expressed in less than the specified “min cells” threshold of cells. |
| [EXPERIMENT]/[Preprocessing] | min genes<br>[type: integer] | Filter cells with less than the specified threshold of “min genes” expressed. |
| [EXPERIMENT]/[Preprocessing] | library size<br>[type: integer] | Library size used for library-size normalization. |
| [EXPERIMENT]/[Preprocessing] | louvain res<br>[type: float] | Louvain clustering resolution (higher resolution means finding more and smaller clusters). |
| [EXPERIMENT]/[Preprocessing] | highly variable number<br>[type: integer] | Number of highly variable genes to identify and keep for the rest of the pipeline. |
| [EXPERIMENT]/[Data] | train<br>[type: file path] | Path to a `.h5ad` file to output the training set. |
| [EXPERIMENT]/[Data] | validation<br>[type: file path] | Path to a `.h5ad` file to output the validation set. |
| [EXPERIMENT]/[Data] | test<br>[type: file path] | Path to a `.h5ad` file to output the test set. |
| [EXPERIMENT]/[Data] | number of genes<br>[type: integer] | Number of genes in the dataset. |

Once the configuration file is ready, run the program with the --preprocess flag, supplying the path to the configuration file with the --config flag.

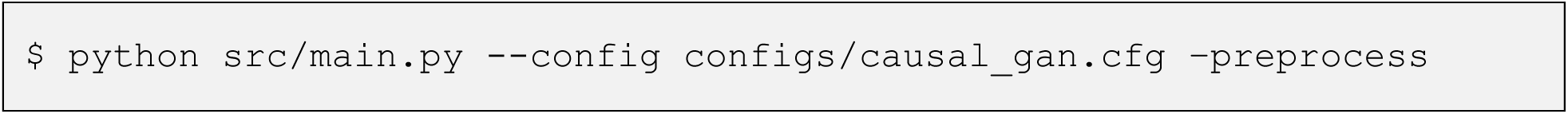

◆ **TROUBLESHOOTING**

Once completed, you will be shown a success message and the preprocessed train, validation, and test sets will be saved as separate .h5ad files in the paths defined under the [Data] section of the configuration file.

*Example of the expected output:*

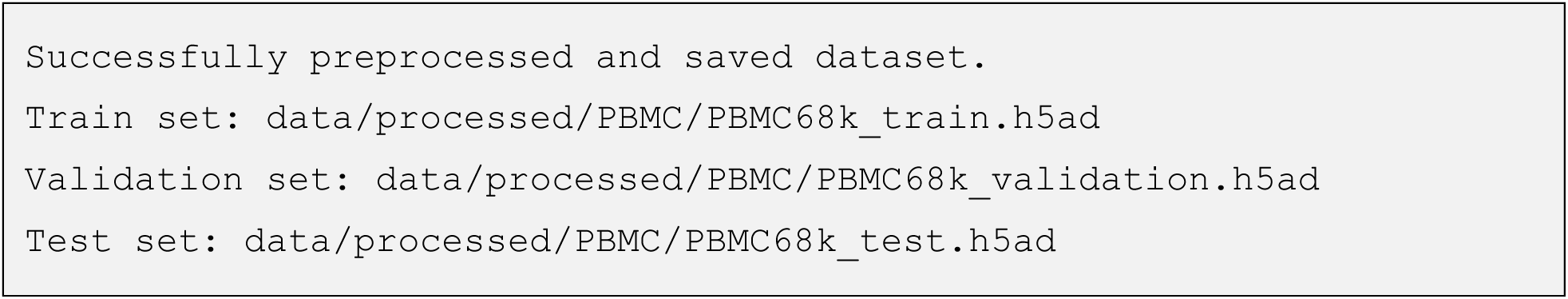

▴ **CRITICAL** Do not replace GRouNdGAN’s preprocessing with a custom preprocessing step. GRouNdGAN specifically requires library-size normalized count data.

### Step 2. GRN Creation

#### A) Imposing a GRN Inferred from Data

##### TIMING ∼8 min for a dataset of around 67000 cells with 4 CPU threads

GRouNdGAN by default uses GRNBoost2[5] to infer a GRN on the preprocessed training set. It then uses the inferred GRN as a scaffold to subsample edges in order to create a causal bipartite directed acyclic graph (DAG). This DAG is then converted and saved as a pickle file in a custom format (see Step 2 - Imposing Custom GRNs) readable to the program. The `–-create_grn` command requires the arguments listed in Table 3, in addition to those previously discussed for Step 1 - Preprocessing. GRouNdGAN provides two ways to subsample the GRN. The first uses `[EXPERIMENT]/[GRN Preparation]/strategy = top`, which includes the top k most important TFs of each gene. The second involves generating two separate GRNs, each containing half of these top k TFs per gene. By setting `[EXPERIMENT]/[GRN Preparation]/strategy = pos ctr` and then `[EXPERIMENT]/[GRN Preparation]/strategy = neg ctr`, the TFs are split based on their parity in the list of each gene’s ranked list of important TFs. Since this alternative method produces two GRNs of identical densities and relatively similar importances, it is useful for comparative and controlled analysis.

**Table 3:** Arguments relevant to the GRN creation command in the configuration file.

| Configuration File Section | Argument | Explanation |
| --- | --- | --- |
| [EXPERIMENT]/[GRN Preparation] | TFs<br>[type: file path] | Path to the file containing TF names (accepts the AnimalTFDB .csv format). |
| [EXPERIMENT]/[GRN Preparation] | k<br>[type: integer] | The number of most important TFs per gene to include in the GRN. |
| [EXPERIMENT]/[GRN Preparation] | strategy<br>[string] | Defines the procedure used to subsample from the inferred GRN. The following options are available:<br>"top": keep the top most important "k" TFs for every gene.<br>"pos ctr": keep the TFs with even parity in the list of top "k" TFs per gene (even indices 0, 2, 4... = top 1, 3, 5, ...).<br>"neg ctr": keep the TFs with odd parity in the list of top "k" TFs per gene (odd indices 1, 3, 5... = top 2, 4, 6, ...). |
| [EXPERIMENT]/ [GRN Preparation] | Inferred GRN<br>[type: file path] | File path to store the GRN inferred by GRNBoost2 (.csv file) |
| [EXPERIMENT]/ [Data] | causal graph<br>[type file path] | File path to store the GRouNdGAN subsampled GRN (.pkl file) |

After completing the relevant sections of the configuration file, execute the following command to run this step:

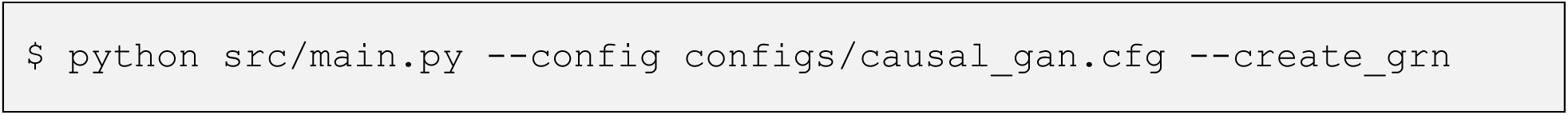

◆ **TROUBLESHOOTING**

Once finished, you will be shown an output detailing the properties of the subsampled GRN and the causal graph will be written in the location specified by [EXPERIMENT]/[Data]/causal graph in the configuration file.

*Example of the expected output:*

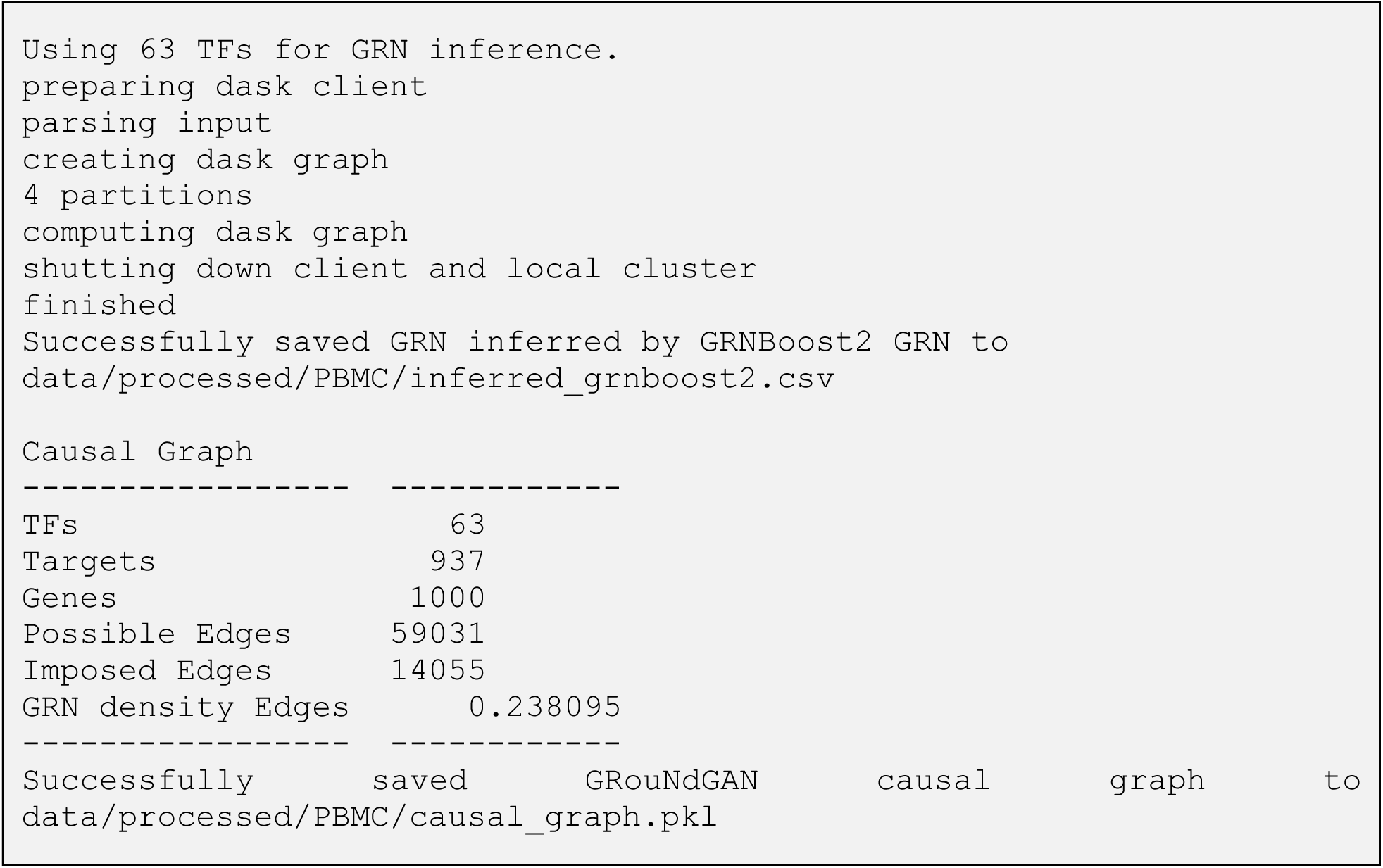

#### B) Imposing a Custom GRN

▲ **CRITICAL STEP** Instead of creating causal graphs through the described subsampling process of a GRNBoost2 inferred GRN, it is possible to impose custom GRNs for greater control over exact edges to include and exclude from the causal graph. Nonetheless, we still advise users to impose causal edges that are biologically meaningful to the reference dataset (through evidence found from literature, computational methods, ChiP-seq datasets, etc.).

▲ **CRITICAL** Since GRouNdGAN’s two-fold objective is to impose a causal structure while generating realistic data points, drastic inconsistencies between the reference data and the imposed causal graph can deteriorate the quality of simulated cells.

GRouNdGAN reads the causal graph from a pickle file storing a python dictionary. In this dictionary, each key corresponds to a gene index, and each value is a Python set containing the indices of its regulating TFs.

Before saving the dictionary, gene and TF names must be converted to their corresponding indices based on their order in the input dataset (.`var_names` attribute in the Anndata file). For example, a dictionary structured with gene/TF names of the form

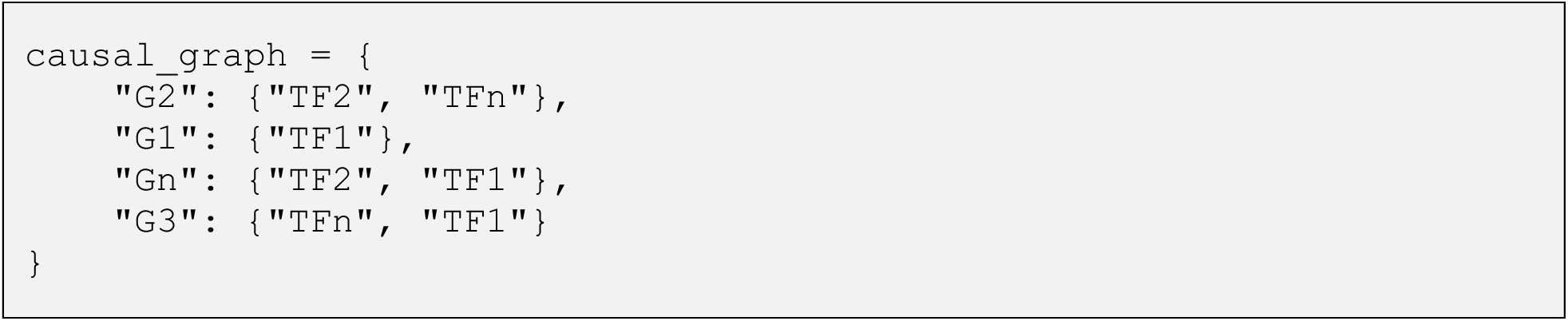

must first be converted into index-based format after resolving each name to its dataset-specific index below:

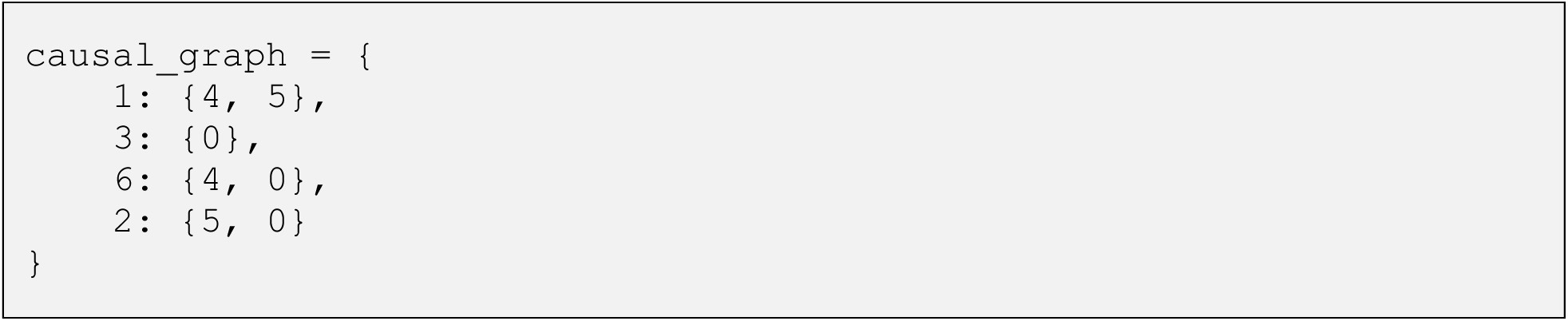

Finally, the causal graph dictionary can be saved as a pickle file in python as shown below.

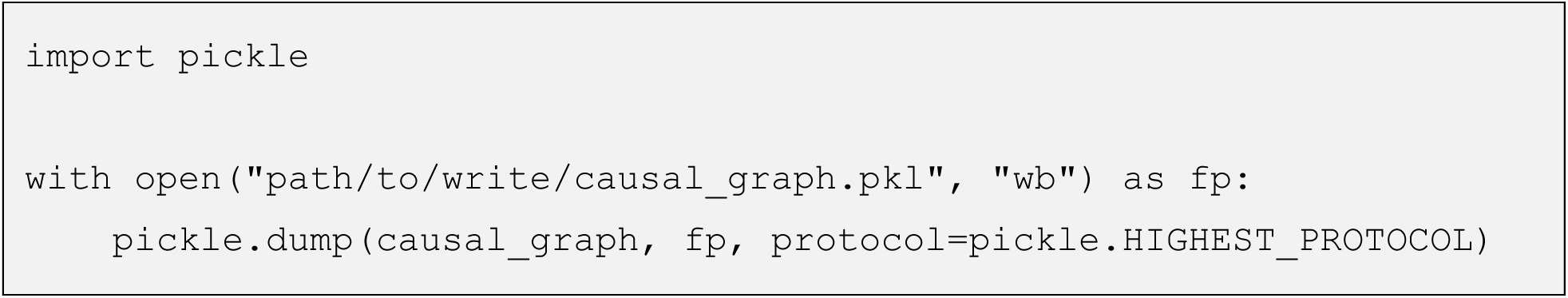

▴ **CRITICAL** If you’re imposing a custom GRN, skip the --create_grn command.

▴ **CRITICAL** The GRN must be a directed bipartite graph. It cannot include cycles.

▲ **CRITICAL** The python dictionary must include all TFs and genes present in the preprocessed dataset. Every target gene must appear as a key in the dictionary, and every TF must appear at least once in the list of values associated with those keys.

▲ **CRITICAL** Make sure the [EXPERIMENT]/[Data]/causal graph] argument in the configuration file points to the newly-created pickle file.

### Step 3. Model Training

#### TIMING ∼75 h on a single NVidia A100SXM4 GPU

When running training command, three folders will be created in the output directory specified by the configuration file (`[EXPERIMENT]/output directory`). The configuration file will also be copied over into this directory.

- `checkpoints/`: contains the `.pth` state dictionaries, including model’s weights, biases, and other parameters.
- `TensorBoard/`: contains stored TensorBoard logs (.tfevent files) for visualizing the model’s training progress and neural graph.
- `TSNE/`: contains t-SNE plots of reference and simulated data.

The frequency at which these directories are updated can be configured through parameters specified in the configuration file, as detailed in Table 4.

**Table 4:** Arguments relevant to the training command in the configuration file.

| Configuration File Section | Argument | Explanation |
| --- | --- | --- |
| `[EXPERIMENT]/[Training]/ [Logging]` | summary frequency<br>[type: integer] | Specifies the frequency at which TensorBoard logs are generated for the GRouNdGAN model. |
| `[EXPERIMENT]/[Training]/ [Logging]` | plot frequency<br>[type: integer] | Specifies the frequency at which t-SNE plots are generated for the GRouNdGAN model. |
| `[EXPERIMENT]/[Training]/ [Logging]` | save frequency<br>[type: integer] | Specifies the frequency at which checkpoints are saved for the GRouNdGAN model. |
| `[EXPERIMENT]/[CC Training]/ [Logging]` | summary frequency<br>[type: integer] | Specifies the frequency at which TensorBoard logs are generated for the causal controller. |
| `[EXPERIMENT]/[CC Training]/ [Logging]` | plot frequency<br>[type: integer] | Specifies the frequency at which t-SNE plots are generated for the causal controller. |
| `[EXPERIMENT]/[CC Training]/ [Logging]` | save frequency<br>[type: integer] | Specifies the frequency at which checkpoints are saved for the causal controller. |

To begin training, use the `--train` flag and along with the path to the configuration file as shown below.

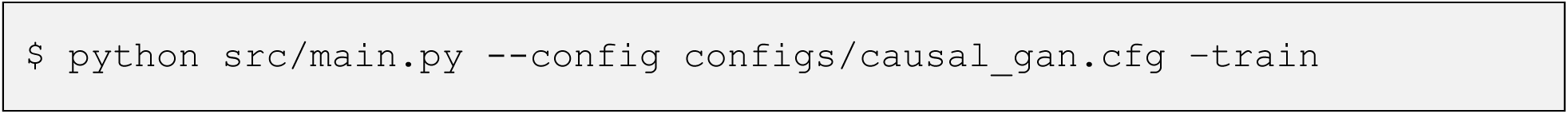

◆ **TROUBLESHOOTING**

*Example of the expected output:*

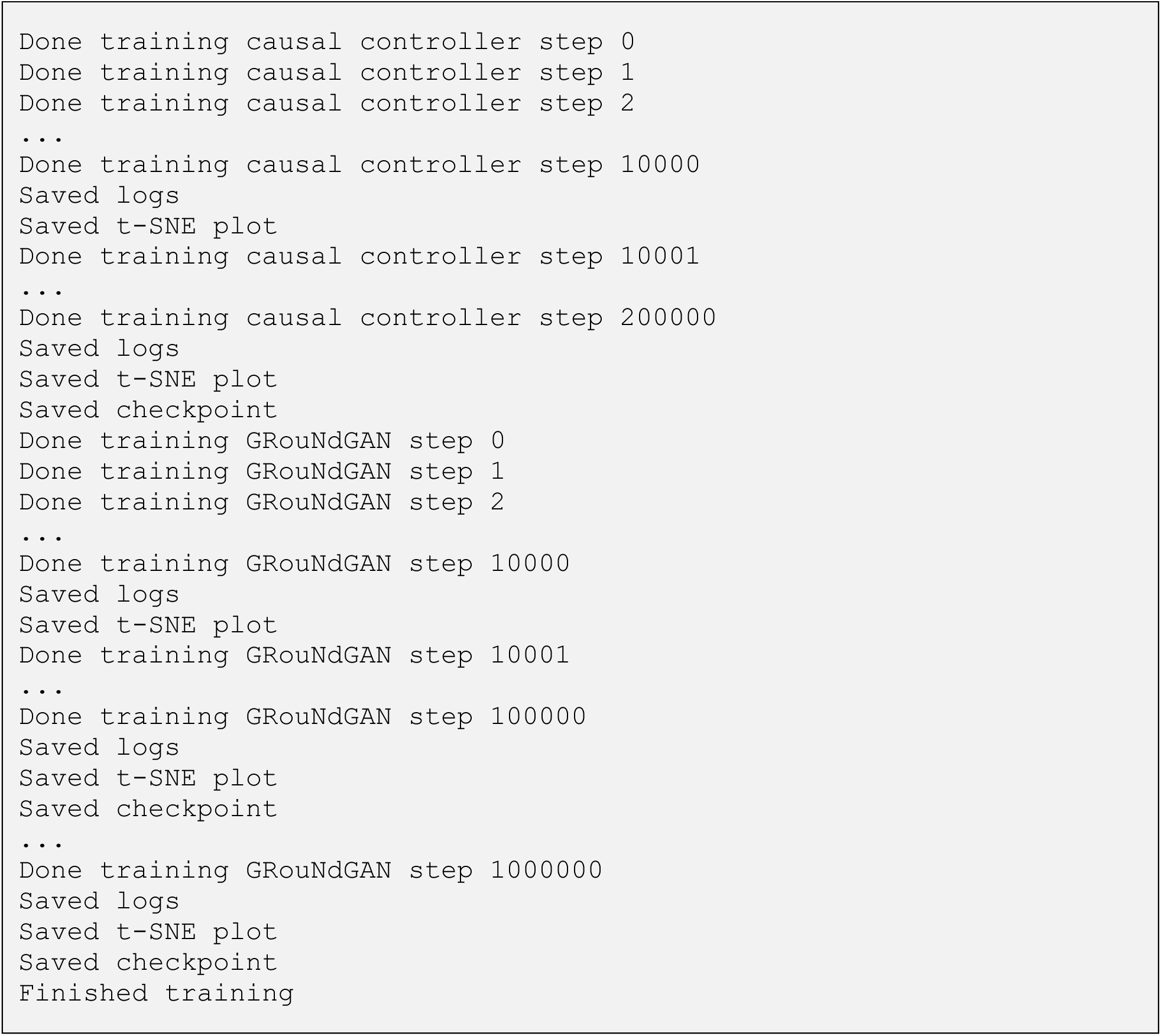

▴ **CRITICAL** While GRouNdGAN supports muti-GPU training, we recommend using a single GPU to minimize overhead. By default, GRouNdGAN trains for 1000000 steps. Changing this setting in the configuration file is not recommended.

▲ **CRITICAL** GRouNdGAN also provides two slurm scripts for training and monitoring on HPC clusters in the scripts/ directory of its code repository.

■ **PAUSE POINT** Users can stop and resume training from a checkpoint by setting [EXPERIMENT]/checkpoint in the configuration file to the latest saved checkpoint (.pth file in [EXPERIMENT]/output directory/checkpoints/) and rerunning the –train command.

Training Monitoring using TensorBoard

TensorBoard is a visualization tool that GRouNdGAN uses to provide its users with an intuitive monitoring and visualization platform. It keeps track of several key scalar values such as generator and critic losses, learning rates, and gradient penalty (Figure 4-b). At regular intervals, it also logs t-SNE plots comparing cells from the validation set with an equal number of simulated cells (Figure 4-a). Furthermore, TensorBoard provides a detailed view of GRouNdGAN’s architecture by rendering the computational graphs of the Wasserstein critic and generator (comprised of causal controller and target generators) at varying levels of complexity as shown in Figure 4-c,d.

**Figure 4:**
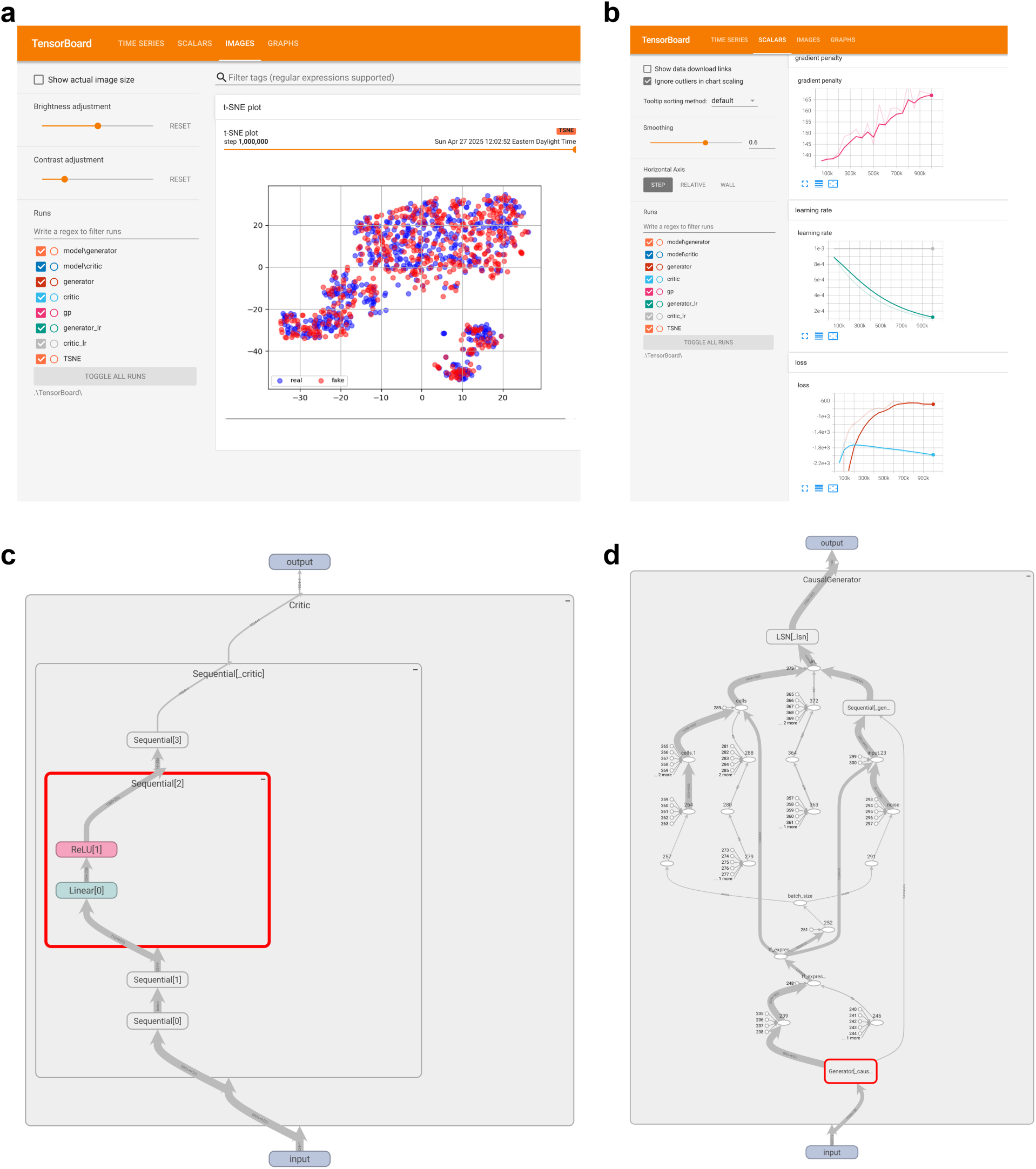
TensorBoard visualizations during GRouNdGAN training. **a** The “images” tab displays a t-SNE plot of reference and generated cells. **b** The “scalars tab” displays key training metrics over time, including gradient penalty, learning rate, and losses for both the critic and generator. **c** and **d** The “graphs” tab visualise the computational graph for the critic and generator networks of GRouNdGAN, showing the model architecture and the flow of gradients during backpropagation.

GRouNdGAN saves TensorBoard event files in [EXPERIMENT]/output directory_CC/Tensorboard for the causal controller training and [EXPERIMENT]/output directory/Tensorboard for the second training step.

Run the following command to start TensorBoard.

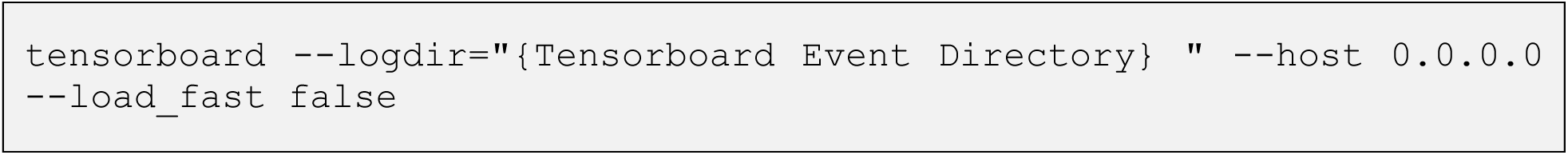

▴ **CRITICAL** Users can replace the {Tensorboard Event Directory} in the above command with a parent directory to compare different training runs of the model.

Once Tensorboard is running, it will output a local URL (by default: http://localhost:6006/) that can be opened in a web browser to access the TensorBoard dashboard. In the case you are running GRouNdGAN on a remote server, you might need to set up port forwarding to access TensorBoard on your local machine. The command below forwards the remote machine’s port 6006 (where TensorBoard is running) to port 6007 on your local machine.

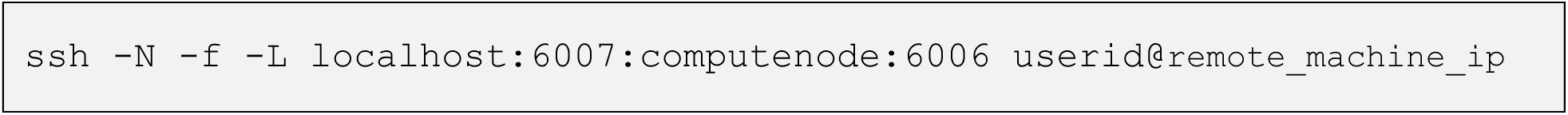

▴ **CRITICAL** The above command might need to be modified depending on your remote server setup, network configuration, and port availability. Ensure that the port on which TensorBoard is running is not blocked by a firewall on the remote server. Some servers might be configured to restrict port forwarding.

### Step 4. Simulation

#### TIMING <30 s

Once training is finished, set the `[EXPERIMENT]/checkpoint` path in the configuration file to the desired model checkpoint (`.pth` file in `[EXPERIMENT]/output directory/checkpoints/`). In most cases, you would want to use the latest checkpoint corresponding to the greatest number of steps. Furthermore, you can control the generation path and number of cells to generate through the configuration file (refer to Table 5).

**Table 5:** Arguments relevant to the generation command in the configuration file.

| Configuration File Section | Argument | Explanation |
| --- | --- | --- |
| `[EXPERIMENT]/ checkpoint` | model checkpoint<br>[type: file path] | Path to the checkpoint from which to load the trained model. |
| `[EXPERIMENT]/[generation]` | number of cells to generate<br>[type: integer] | Number of cells to simulate. |
| `[EXPERIMENT]/[generation]` | generation path<br>[type: directory, Optional] | Path where to save the simulated dataset (`simulated.h5ad`). If left undefined, `simulated.h5ad` will be saved to `[Experiment]/output directory/`. |

Running the following command will save the generated cells as a .h5ad file (simulated.h5ad).

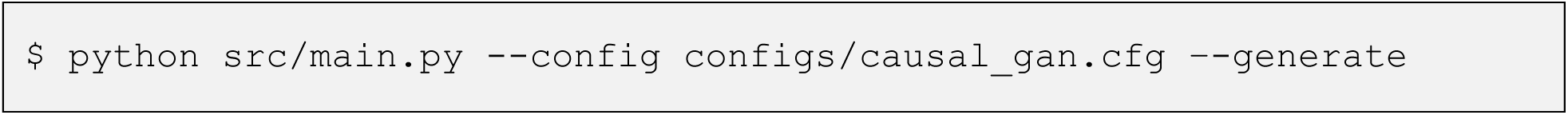

*Example of the expected output:*

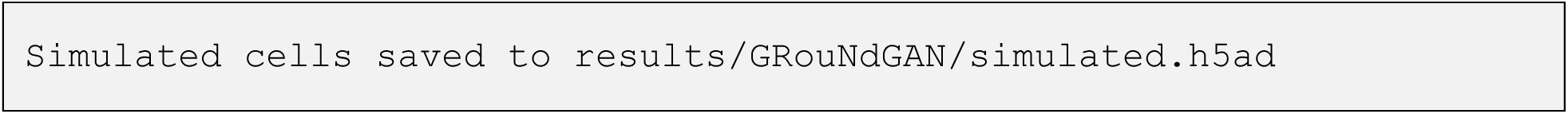

■ **PAUSE POINT** Remaining steps are optional. Users can stop here and use the saved simulated dataset for benchmarking, data augmentation, and other external analyses.

Step 5. Assessing simulated data quality (Optional)

##### TIMING <30 s

This step provides a simple command for assessing the quality of simulated scRNA-seq data compared to its reference experimental dataset using the following qualitative and quantitative metrics:

- t-SNE plot visualization of jointly embedded experimental and simulated cells. Greater overlap indicates better alignment.
- Euclidean distance between the mean expression profiles of real and simulated cells. A lower value indicates higher similarity.
- Cosine distance between the mean expression profiles of real and simulated cells. A lower value indicates higher directional similarity and better alignment.
- AUROC of a random forest classifier trained to distinguish simulated cells from experimental cells. An AUROC closer to 0.5 indicates that the classifier fails to tell the two classes of cells apart, implying more realistic simulation.
- Maximum Mean Discrepancy (MMD) to quantify distributional similarity of experimental and simulated cells. A lower value is desired.
- Mean integration local inverse Simpson’s index (miLISI) to measure local mixing of real and simulated cells in the t-SNE space. A value closer to 2 indicates good integration and mixing.

All metrics are computed by comparing the test set to an equal number of cells sampled from the simulated data. As a control, we also compute the Euclidean distance, Cosine distance, MMD, and miLISI between two halves of the reference test set to establish a baseline for comparison.

You can specify which evaluation metrics to compute by setting the corresponding flags in the configuration file (see Table 6). Additionally, you can use GRouNdGAN’s evaluation framework to assess simulated datasets generated by external simulators by updating `[EXPERIMENT]/ [Evaluation]/simulated data path` in the configuration file to point to the desired dataset.

**Table 6:**
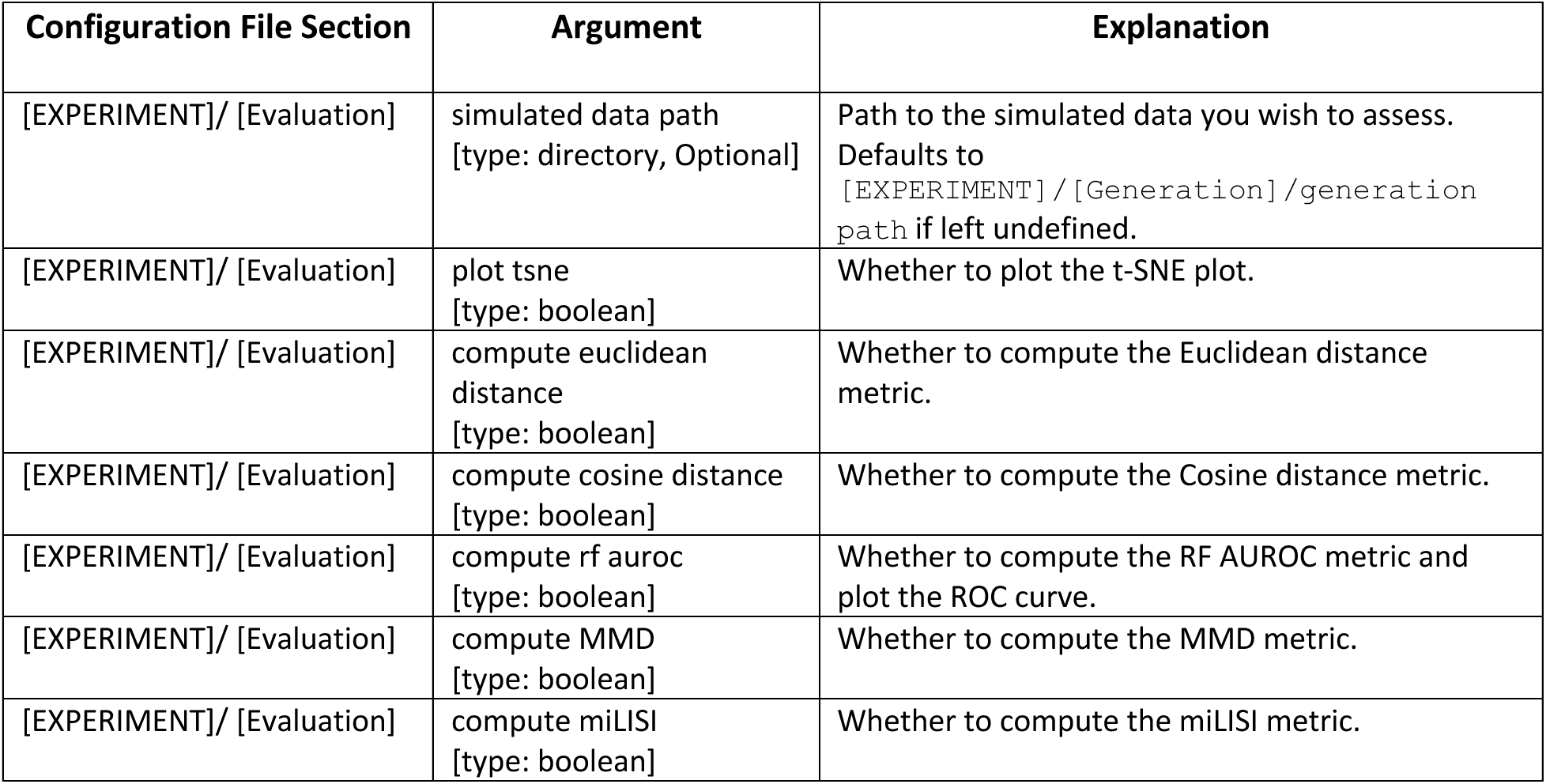
Arguments relevant to the evaluation command in the configuration file.

To run the evaluation pipeline with the specified configuration, execute the following command.

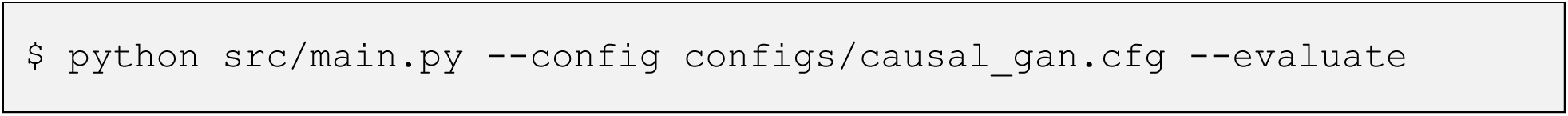

Output plots (following the format of Figure 5-a,b) will be saved to [EXPERIMENT]/output directory/.

**Figure 5:**
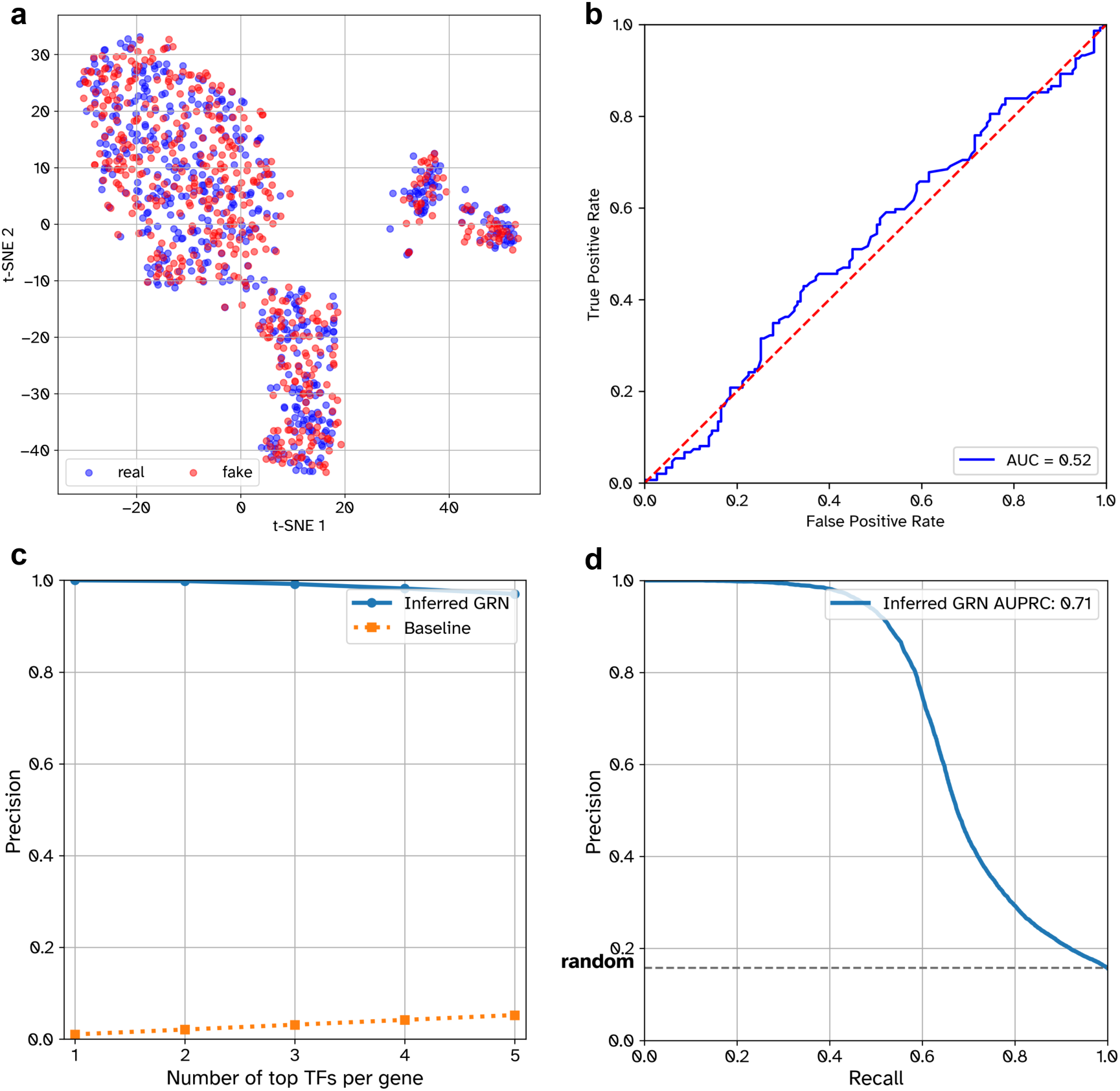
Example of expected plot outputs saved by the evaluation and benchmarking commands. **a** t-SNE plot of simulated and test set experimental cells. **b** Receiver Operating Characteristic (ROC) curve of a random forest classifier distinguishing between simulated cells and test set experimental cells. **c** Precision at k plot showing the proportion of true regulators among the top-k predicted transcription factors for each gene. **d** Precision-Recall (PR) curve illustrating global GRN reconstruction performance. The baseline corresponds to the performance of a random predictor which also corresponds to the density of the ground truth GRN.

*Example of the expected output:*

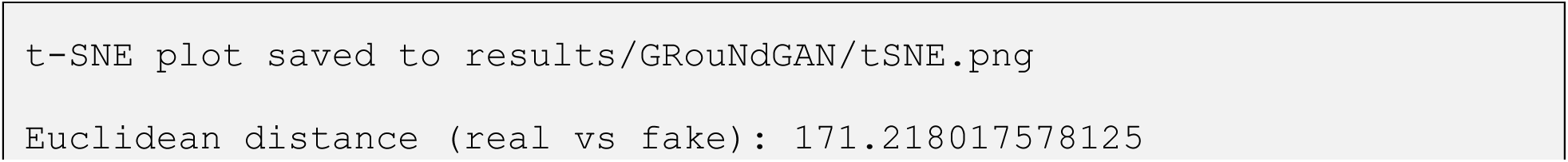
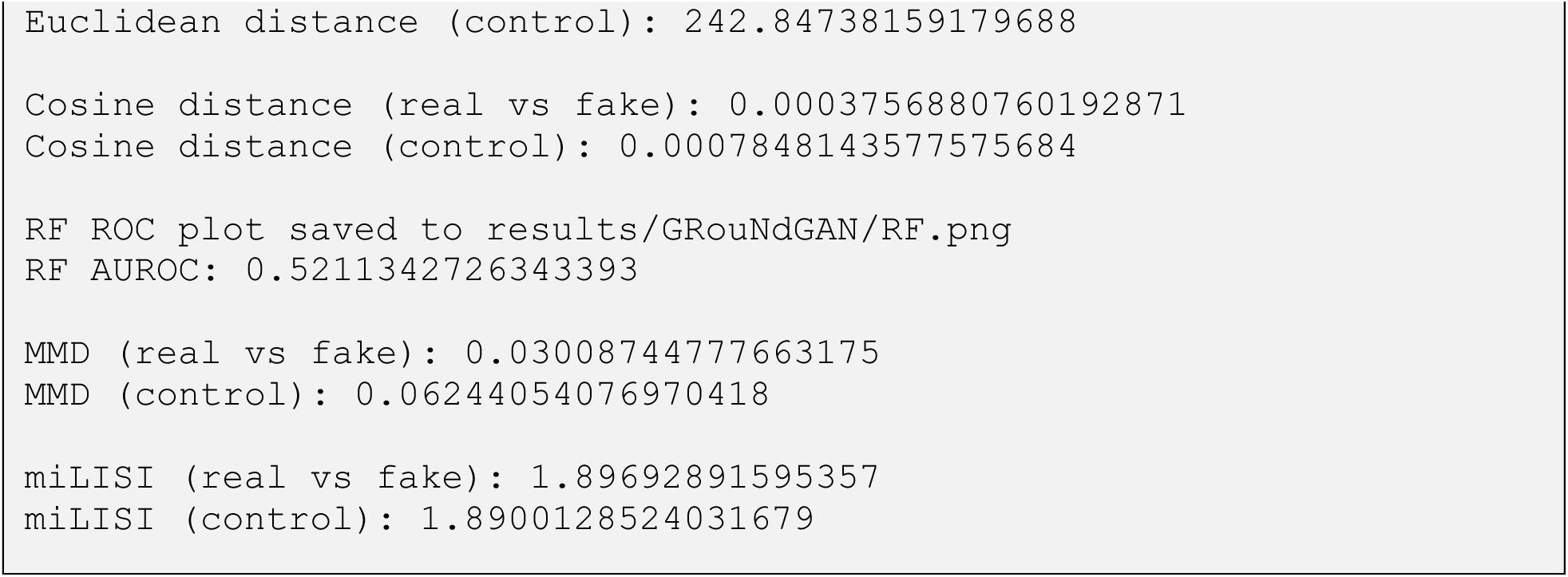

### Step 6. GRN inference benchmarking (Optional)

#### TIMING <20 s

One of GRouNdGAN’s main goals is to provide tools and ground truth for benchmarking GRN inference methods. This command evaluates how faithful GRNs inferred on GRouNdGAN-generated data are to the one imposed onto the model and used for simulation. To run this command, you must provide the path to the inferred GRN through the configuration file using the `[EXPERIMENT]/[GRN Benchmarking]/grn to benchmark]` field (refer to Table 7). The GRN should be a tab-separated `.csv` file with three columns in the following order:

1. TF
2. Gene
3. Importance

**Table 7:** Arguments relevant to the benchmarking command in the configuration file.

| Configuration File Section | Argument | Explanation |
| --- | --- | --- |
| [EXPERIMENT] / [GRN Benchmarking] | grn to benchmark<br>[type: file path] | Path to the inferred GRN you wish to benchmark. |
| [EXPERIMENT] / [GRN Benchmarking] | ground truth save path<br>[type: directory, Optional] | If specified, the ground truth GRN (GRouNdGAN's causal graph) will be exported to the provided path. |
| [EXPERIMENT] / [GRN Benchmarking] | plots save path<br>[type: path] | Folder in which to save generated plots. |
| [EXPERIMENT] / [GRN Benchmarking] | compute precision at k<br>[type: boolean] | Whether to compute precision at k. |
| [EXPERIMENT] / [GRN Benchmarking] | k<br>[type: integer] | How many of the top k regulators to consider when computing precision at k. |
| [EXPERIMENT] / [GRN Benchmarking] | compute pr<br>[type: boolean] | Whether to plot the PR curve and compute AUPRC of GRN recovery. |

Headers are optional; only the column order matters.

*Example format:*

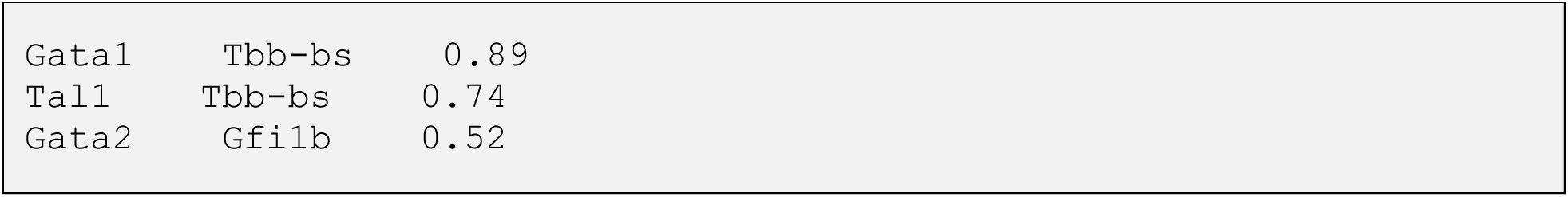

When enabled in the configuration file, the GRN benchmarking command generates one or both of the following plots, which are saved to the directory specified in [EXPERIMENT]/[GRN Benchmarking]/plots save path:

- Precision at k: Evaluates the proportion of the top-k inferred TFs for each gene are true regulators. The value of k is configurable.
- Precision-Recall (PR) curve: Computed across all predicted edges in the inferred GRN.

Run this step using the command below.

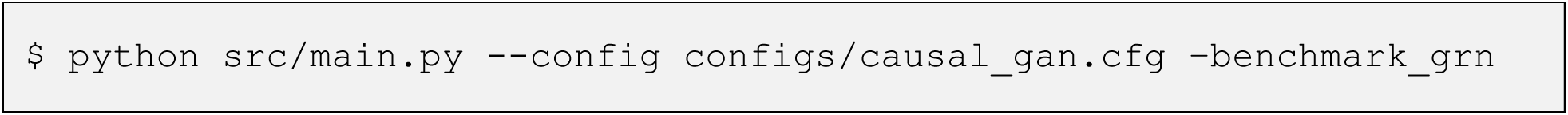

Both plots (see Figure 5-c, d) are saved as .png files, and relevant evaluation metrics such as precision at k scores and the area under the precision recall curve (AUPRC) are printed to the console.

*Example of the expected output:*

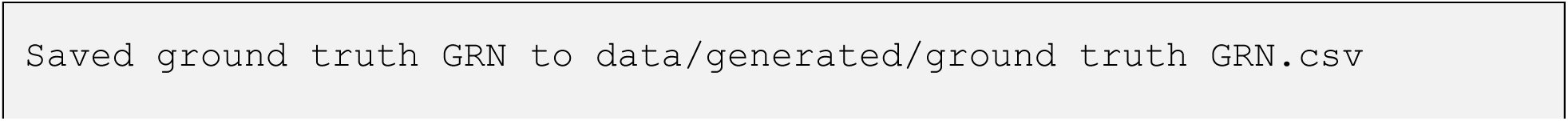
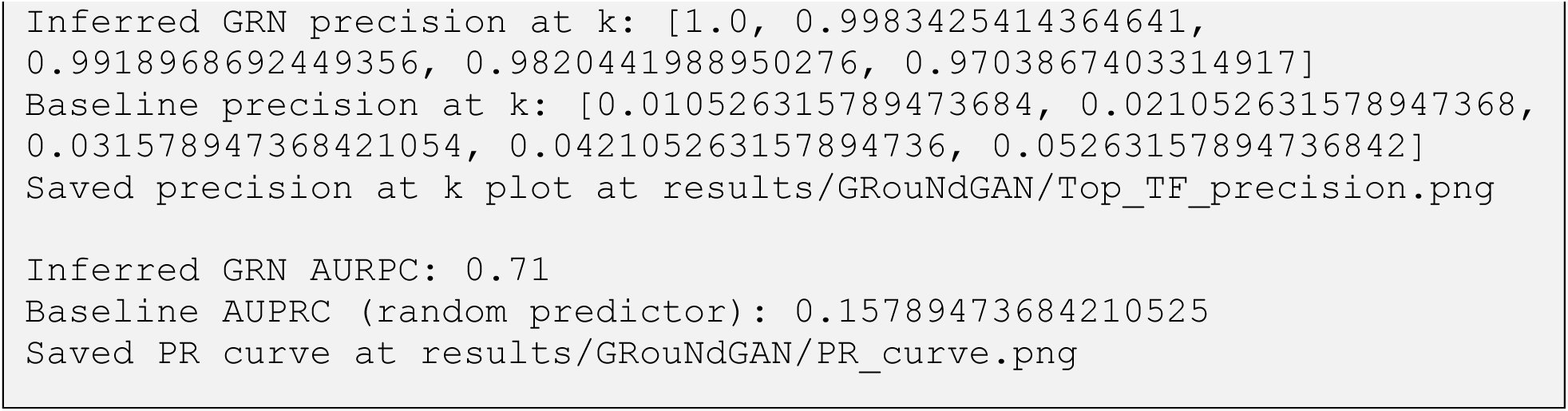

■ **PAUSE POINT** The ground truth GRN can be exported as a `.csv` file to be used for additional analysis or use in external benchmarking tools. Simply specify `[EXPERIMENT]/[GRN Benchmarking]/ground truth save path` in the configuration file before running the command.

Step 7. Perturbation prediction (Optional)

- ● TIMING <1 min

Once trained, GRouNdGAN can sample from interventional distributions and perform in silico perturbation experiments at inference time. It enables the creation of matched case/control comparison as perturbations are applied to the same batch of cells (batch size of 1024 by default). To simulate a perturbation, provide a list of TFs to perturb using `[EXPERIMENT]/[Perturbation]/tfs to perturb`, along with the corresponding target values in `[EXPERIMENT]/[Perturbation]/perturbation values`. Each TF is matched to a value by its position in the list, with entries separated by spaces (as shown in Table 8).

**Table 8:**
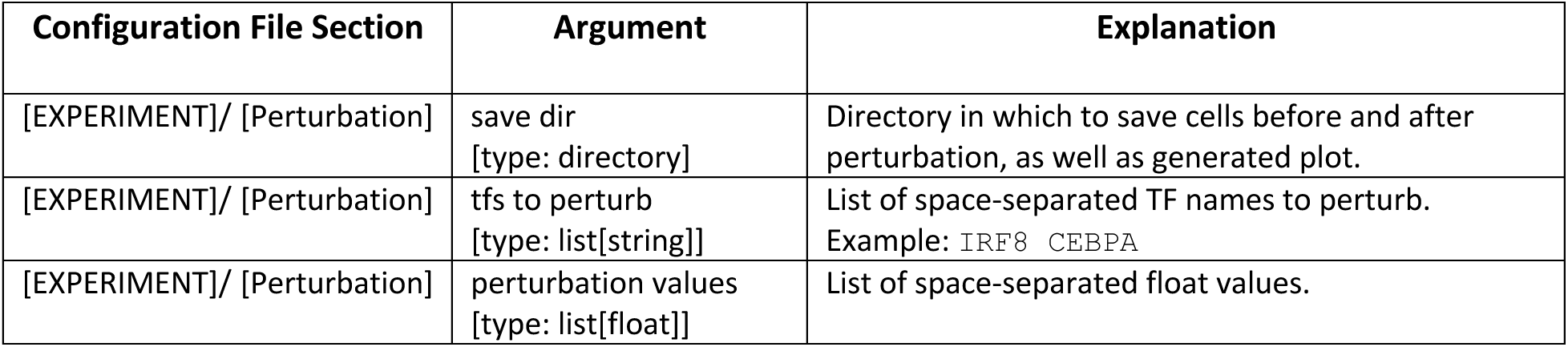
Arguments relevant to the perturbation command in the configuration file.

▴ **CRITICAL** Ensure all TFs specified in [EXPERIMENT]/[Perturbation]/tfs to perturb exist in the causal graph and that their count matches the number of entries in [EXPERIMENT]/[Perturbation]/perturbation.

Once the experiment parameters are defined, you can run the perturbation experiment with the following command.

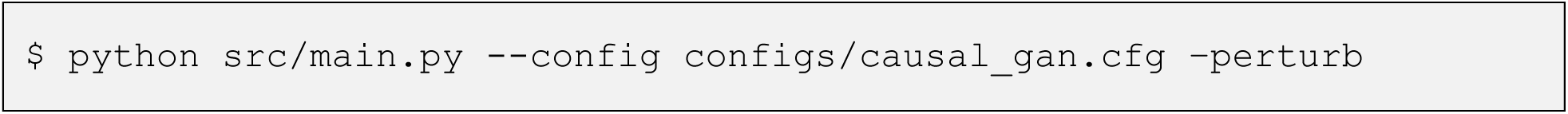

◆ **TROUBLESHOOTING**

The cell states before and after perturbation will be saved as `before_perturbation.h5ad` and `after_perturbation.h5ad`, respectively, in the specified output directory. There is a one-to-one correspondence between cells in the two simulated datasets (row and column indices match). These files can be used for downstream or secondary analysis. In addition to the data files, a series of plots, similar to Figure 6, will be generated and saved to visualize the effects of the perturbation.

**Figure 6:**
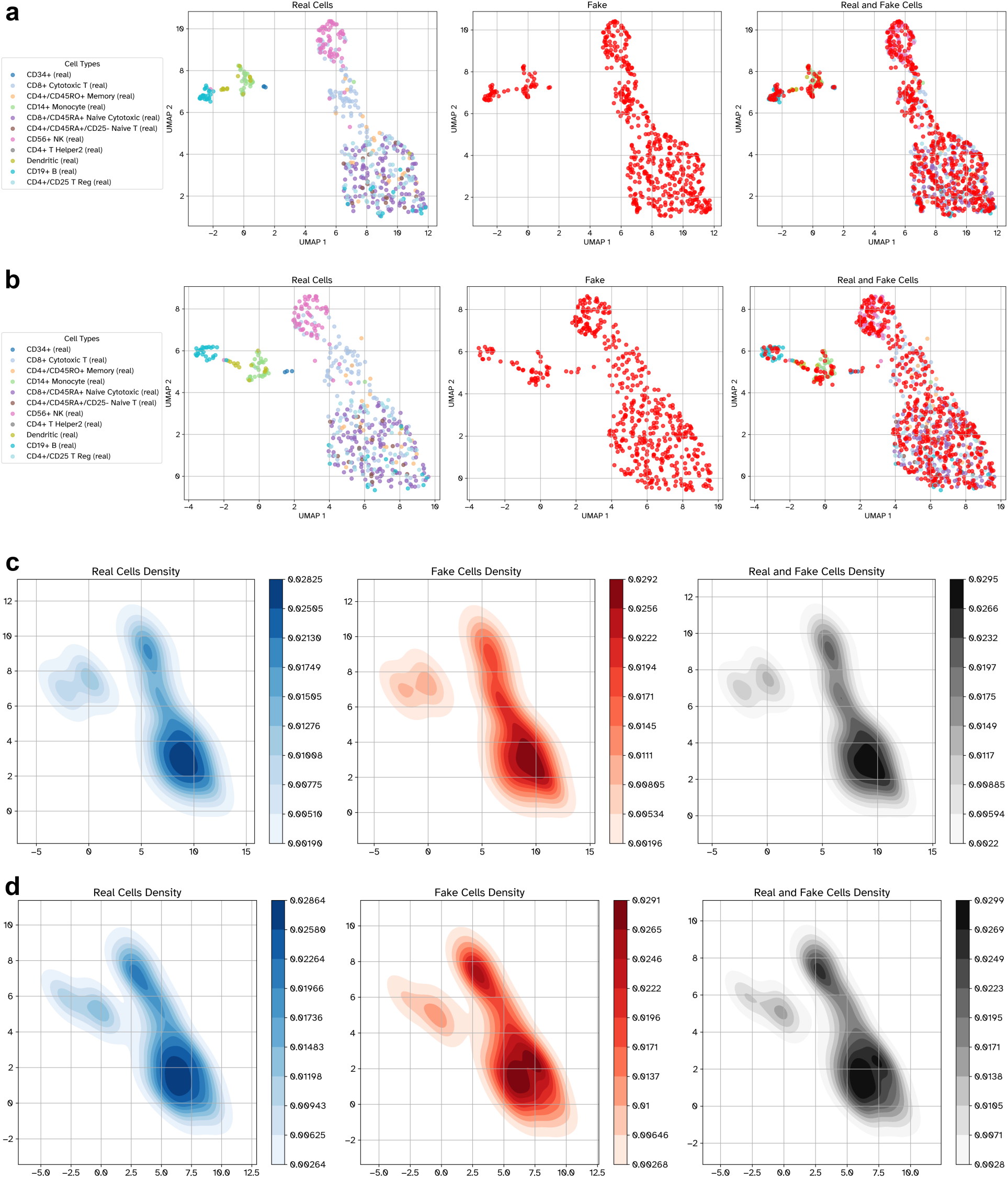
Example of expected plot outputs saved by the perturbation command. **a** UMAP embedding of experimental cells (left), unperturbed simulated cells (middle), and overlayed on top of each other (right). **b** similar to **a** but simulated cells were generated under perturbation. **c** Cell density in the UMAP embedding for experimental cells (left), unperturbed simulated cells (middle), and together (right). **d** similar to **c** but simulated cells are perturbed.

*Example of the expected output:*

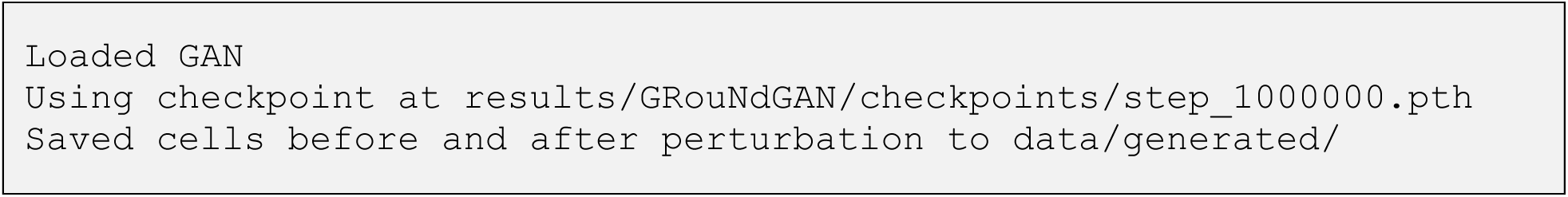

## Timing

The reported execution times correspond to running GRouNdGAN on the PBMC dataset, which comprises 67579 cells, using the hardware configuration detailed in the Equipment section. The duration of step 1 (Preprocessing) and step 2 (GRN creation) primarily scale with the dataset size. In contrast, the execution of step 3 (Model training) is largely dependent on the density of the imposed GRN and GPU specification. The remaining steps exhibit comparatively lower sensitivity to these factors. Installation times are predominantly affected by download speed and server loads.

Software installation - local: ∼12 min

Software installation - running through the Docker image: ∼6 min Software installation - building Docker image from Dockerfile: ∼13 min Software installation - using Singularity: ∼7 min

Procedure - Step 1, Preprocessing: <1 min Procedure - Step 2, GRN creation: ∼8 min

Procedure - Step 3, Model training: ∼75 h Procedure - Step 4, Simulation: <30 s

Procedure - Step 5, Data quality evaluation: <30 s Procedure - Step 6, GRN inference benchmarking: <20 s Procedure - Step 7: Perturbation prediction: <1 min

Procedure - End to end (Steps 1-7): ∼75 h, 12 min

### General notes and troubleshooting

Table 9 contains common issues and how to troubleshoot them.

**Table 9:** Troubleshooting table.

| Step | Problem | Possible reason | Solution |
| --- | --- | --- | --- |
| 1, 2 | Error message:<br>`check_values_indices_shape_match` | Mismatch between the number of genes in train, validation, or test sets and configuration parameters or GRN.<br>The `.X` attribute in the `anndata` input is a sparse expression matrix. | Verify `[EXPERIMENT]/[Preprocessing]/ highly variable number` and `[EXPERIMENT]/[Data]/number of genes` parameters in the configuration file against the input dataset.<br>If the input expression matrix is sparse, convert it into a dense matrix using `adata.X.toarray()`. |
| 3 | Error message:<br>`torch.cuda.OutOfMemoryError: CUDA out of memory` | Insufficient video memory on GPU. | Switching to a GPU with a larger VRAM.<br>Making the input GRN sparser. If the `-create_grn` command is used to construct the input GRN, making the hyperparameter `[GRN Preparation]/k` in the configuration file smaller will result in a sparser GRN. Reducing GRN size might result in a lower quality simulation.<br>Reducing the `[EXPERIMENT]/[Model]/depth per gene` and `[EXPERIMENT]/[Model]/width per gene` hyperparameters in the configuration file to 2 and 1, respectively. This will make GRouNdGAN's neural network smaller to fit in a smaller VRAM. This option might result in a lower quality simulation. |
| 3 | Error message:<br>`what(): mat1 and mat2 shapes cannot be multiplied` | The specified number of genes is inconsistent with the GRN. | Ensure that the number of nodes in the GRN correspond to the `[EXPERIMENT]/[Data]/number of genes` parameter in the configuration file. |
| 7 | Error message:<br>`ValueError: 'TF' is not in list` | The TF targeted for perturbation does not exist in the GRN. | Ensure that all the TFs listed in the `[EXPERIMENT]/[Perturbation]/tfs to perturb` parameter of the configuration file exist in the dataset and are regulators (not targets) in the GRN. |

## Expected results

Running GRouNdGAN’s GRN-guided simulation procedure (Steps 1 to 4) produces a single final output: a `simulated.h5ad` file containing simulated scRNA-seq data. The file preserves the original gene annotations from the input data and assigns sequential labels to generated cells (e.g., Cell1, Cell2, …). The simulated expression matrix is stored in the `.X` attribute, where rows correspond to cells and columns to genes. Furthermore, `.var` and `.obs` store gene and cell metadata, respectively. This structured output format ensures compatibility with downstream python and R single-cell analysis workflows. A collection of such output datasets paired with the GRN used for simulation can be found on GRouNdGAN’s website at https://emad-combine-lab.github.io/GRouNdGAN/benchmarking.html.

The perturbation prediction procedure generates two .h5ad files: `before_perturbation.h5ad` and `after_perturbation.h5ad`. They contain expression matrices (in the `.X` attribute) of identical dimensions, representing cellular states before and after perturbation, respectively. Both include the same set of genes and annotations. Additionally, cells are matched one-to-one between the two files, meaning that each cell (e.g., “Cell 1”) in `before_perturbation.h5ad` corresponds directly to its perturbed counterpart in `after_perturbation.h5ad`.

Given a reasonably sized training set and a sufficiently representative GRN, we expect the simulated output data to be realistic (i.e., mimicking experimental training data) and true to the imposed GRN. Under the same conditions, we also expect that the perturbation results will align with biological expectations, though perturbation prediction relies more heavily on the quality of the input GRN. We provide guidelines for evaluating the quality of the output data using five metrics (see the Procedure – Step 5) and comparing it to a control baseline, where the control is derived from the data itself. Furthermore, it is expected that the generator and critic losses, which can be monitored using TensorBoard (see Procedure – Step3), will decrease as the model undergoes training, showing better performance over time.

## Supplementary Information

The online version contains supplementary material available in Supplementary File S1.

### Authors’ contributions

All authors contributed to the design of the protocol, data analyses, and writing the manuscript.

A.E. supervised the study. Y.Z. implemented the protocol, conducted computational analyses, developed the website, and managed releases across various formats.

### Funding

This work was supported by a Discovery grant from Natural Sciences and Engineering Research Council of Canada (NSERC) [RGPIN-2019-04460] (AE), a NOVA grant funded by NSERC and Fonds de recherche du Québec – Nature et technologies (FRQNT) [2024-NOVA-344140], and by a grant from Canada Foundation for Innovation (CFI) JELF [project 40781] (AE). This work was enabled in part by support provided by Calcul Québec (www.calculquebec.ca) and the Digital Research Alliance of Canada (alliancecan.ca)

### Data Availability

The dataset used to demonstrate the protocol is publicly available. The PBMC-ALL scRNA-seq dataset (healthy donor A) can be downloaded through the 10x Genomics website at https://support.10xgenomics.com/single-cell-gene-expression/datasets/1.1.0/fresh_68k_pbmc_donor_a?. The GRouNdGAN-simulated PBMC-CTL dataset (100k cells by 1000 genes,including TFs), along with the imposed ground truth GRN, is available on GRouNdGAN’s website at https://emad-combine-lab.github.io/GRouNdGAN/benchmarking.html#pbmc-ctl.

### Code Availability

An implementation of GRouNdGAN is freely available under the terms of the GNU Affero General Public License v3.0 on GitHub at https://github.com/Emad-COMBINE-lab/GRouNdGAN. The software is written in Python 3.9.6 using the PyTorch deep learning framework (v. 1.13.1). To ensure ease of use and reproducibility, a pre-build container image is also available on DockerHub (https://hub.docker.com/r/yazdanz/groundgan) and an archived version can be found on Zenodo (https://doi.org/10.5281/zenodo.11068245). Visit the GRouNdGAN website (https://emad-combine-lab.github.io/GRouNdGAN) to explore an expanding collection of readily available simulated datasets for benchmarking GRN inference algorithms, along with the latest updates on running the protocol.

