## Supplementary File S1 for "Causal single-cell RNA-seq simulation, in silico perturbation, and GRN inference benchmarking using GRouNdGAN-Toolkit"

### Supplementary Notes

#### Default GRouNdGAN configuration file.

```
[EXPERIMENT]
output directory = results/GRouNdGAN
device = cuda ; we will let the program choose what is available
checkpoint  ; set value to use a trained model

    [Preprocessing]
    10x = True
    raw = data/raw/PBMC/
    validation set size = 1000
    test set size = 1000
    annotations = data/raw/PBMC/barcodes_annotations.tsv
    min cells = 3 ; genes expressed in less than 3 cells are discarded
    min genes = 10 ; cells with less than 10 genes expressed are discarded
    library size = 20000 ; library size used for library-size normalization
    louvain res = 0.15 ; Louvain clustering resolution (higher resolution means finding more and
smaller clusters)
    highly variable number = 1000 ; number of highly variable genes to identify

    [GRN Preparation]
    TFs = data/raw/Homo_sapiens_TF.csv
    k = 15 ; k is the number of top most important TFs per gene to include in the GRN
    Inferred GRN = data/processed/PBMC/inferred_grnboost2.csv

    ; "top" for selecting top k edges
    ; "pos ctr" for generating positive control GRNs (even indices 0, 2, 4... = top 1, 3, 5, ...)
    ; "neg ctr" for generating negative control GRNs (odd indices 1, 3, 5... = top 2, 4, 6, ...)
    ; note that k has to be a pair number for strategy=ctr
    strategy = top

    [Data]
    train = data/processed/PBMC/PBMC68k_train.h5ad
    validation = data/processed/PBMC/PBMC68k_validation.h5ad
    test = data/processed/PBMC/PBMC68k_test.h5ad
    number of genes = 1000

    ; this causal graph is a pickled nested dictionary
    ; nested dictionary keys are gene indices
    ; the dictionary is of this form:
    ; {381: {51, 65, 353, 664, 699},
    ; 16: {21, 65, 353, 605, 699},
    ; ...
    ; 565: {18, 51, 65, 552, 650}}
    ; In this example, 381, 16, and 565 are gene indices in the input dataset
    ; Each key's (gene's) value is the indiced of its regulating TFs in the input dataset
    ; A tutorial will be made available in the future.

    causal graph = data/processed/PBMC/causal_graph.pkl

    [Generation]
    number of cells to generate = 10000
    generation path ; will save to [Experiment]/output directory/simulated.h5ad if left undefined

    [Evaluation]
    simulated data path ; will use [Generation]/generation path if left undefined
    plot tsne = True ; Note: has to be true in order to run miLISI
    compute euclidean distance = True
    compute cosine distance = True
    compute rf auroc = True
    compute MMD = True
    compute miLISI = True ; plot tsne has to be True for this to work

    [GRN Benchmarking]
    grn to benchmark = path/to/inferred/grn.csv
    ground truth save path = data/generated/
    plots save path = results/GRouNdGAN/
```

```
    compute precision at k = False
    k = 5
    compute pr = True

    [Perturbation]
    save dir = data/generated/

    ; tfs to perturb and perturbation values are paired. The two lists need to be of the same
size.
    tfs to perturb ;= TF names separated by a space (ex: IRF8 CEBPA)
    perturbation values ;= Float values separated by a space to set respecive TF in tfs to
perturb (ex: 0 100.2)

    [Model]
    type = causal GAN
    noise per gene = 1
    depth per gene = 3
    width per gene = 2
    critic layers = 1024 512 256
    labeler layers = 2000 2000 2000
    latent dim = 128 ; noise vector dimensions
    lambda = 10 ; regularization hyper-parameter for gradient penalty


    [Training]
    batch size = 1024
    critic iterations = 5 ; iterations to train the critic for each iteration of the generator
    maximum steps = 1000000
    labeler and antilabeler training intervals = 1

        [Optimizer]
        ; coefficients used for computing running averages of gradient and its square
        beta1 = 0.5
        beta2 = 0.9

        [Learning Rate]
        generator initial = 0.001
        generator final = 0.0001
        critic initial = 0.001
        critic final = 0.001
        labeler = 0.0001
        antilabeler = 0.0001


        [Logging]
        summary frequency = 10000
        plot frequency = 10000
        save frequency = 100000

    [CC Model]
    type = GAN ; Non-conditional single-cell RNA-seq GAN
    generator layers = 256 512 1024
    critic layers = 1024 512 256
    latent dim = 128 ; noise vector dimensions
    lambda = 10 ; regularization hyper-parameter for gradient penalty


    [CC Training]
    batch size = 128
    critic iterations = 5 ; iterations to train the critic for each iteration of the generator
    maximum steps = 200000

        [CC Optimizer]
        ; coefficients used for computing running averages of gradient and its square
        beta1 = 0.5
        beta2 = 0.9

        [CC Learning Rate]
        generator initial = 0.0001
        generator final = 0.00001
        critic initial = 0.0001
```

```
critic final = 0.00001

[CC Logging]
summary frequency = 10000
plot frequency = 10000
save frequency = 100000
```

#### Supplementary Tables

**Table S1:** Number of cells in the train and test set of cell type-specific GRouNdGAN models. Relative abundance values reflect the proportion of each cell type in the PBMC dataset.

| Cell Type | Train Set Size | Test Set Size | Relative Abundance (%) |
|---|---|---|---|
| CD8+ Cytotoxic T | 19773 | 1000 | 30.29 |
| CD8+/CD45RA+/Naïve Cytotoxic | 15666 | 1000 | 24.30 |
| CD56+ NK | 7776 | 1000 | 12.80 |
| CD4+ /CD25 T Reg | 5187 | 1000 | 9.02 |
| CD19+ B | 4908 | 1000 | 8.61 |

**Table S2:** Performance of GRouNdGAN in generating realistic scRNA-seq data using an alternative cell-type-specific GRN setup using the PBMC dataset. In this setting, the dataset is first partitioned by cell type, followed by independent preprocessing and GRN construction for each subset. The metrics are calculated by comparing a held-out test set of 1000 real cells and a simulated dataset of the same size. In the imposed GRN each gene is regulated by a maximum of 15 TFs. The default algorithm GRNBoost2 was used with the training experimental data to form the imposed GRN. For the cosine distance, Euclidean distance, and MMD, values closer to zero are desired. For RF AUROC a value closer to 0.5 and for miLISI a value closer to 2 reflect a more realistic simulation.

| Simulator | Cell Type | Cosine distance | Euclidean distance | MMD | RF AUROC | miLISI |
|---|---|---|---|---|---|---|
| GRouNdGAN | CD8+ Cytotoxic T | 0.00025 | 140 | 0.026 | 0.49 | 1.905 |
| Control | CD8+ Cytotoxic T | 0.00041 | 160 | 0.027 | 0.5 | 1.912 |
| GRouNdGAN | CD8+/CD45RA+/Naïve Cytotoxic | 0.00067 | 163 | 0.029 | 0.52 | 1.895 |
| Control | CD8+/CD45RA+/Naïve Cytotoxic | 0.00034 | 116 | 0.026 | 0.5 | 1.900 |
| GRouNdGAN | CD56+ NK | 0.0010 | 195 | 0.031 | 0.54 | 1.918 |
| Control | CD56+ NK | 0.00047 | 133 | 0.026 | 0.5 | 1.907 |
| GRouNdGAN | CD4+ /CD25 T Reg | 0.00020 | 93 | 0.029 | 0.55 | 1.896 |
| Control | CD4+ /CD25 T Reg | 0.00023 | 109 | 0.025 | 0.5 | 1.919 |
| GRouNdGAN | CD19+ B | 0.00032 | 116 | 0.027 | 0.56 | 1.910 |
| Control | CD19+ B | 0.00061 | 165 | 0.024 | 0.5 | 1.909 |
| GRouNdGAN | PBMC-ALL | 0.00020 | 151 | 0.014 | 0.51 | 1.900 |
